# MIXL1 Activation in Endoderm Differentiation of Human Induced Pluripotent Stem Cells

**DOI:** 10.1101/2024.03.06.583475

**Authors:** Pierre Osteil, Sarah Withey, Nicole Santucci, Nader Aryamanesh, Ignatius Pang, Nazmus Salehin, Jane Sun, Annie Qin, Jiayi Su, Hilary Knowles, Xiucheng Bella Li, Simon Cai, Ernst Wolvetang, Patrick P. L. Tam

## Abstract

Human induced pluripotent stem cells (hiPSC) possess the ability to differentiate into a multitude of cell and tissue types but display heterogeneous propensity of differentiation into specific lineage. Characterization of the transcriptome of eleven hiPSC lines showed that activation of *MIXL1* at the early stage of stem cell differentiation correlated with higher efficacy in generating definitive endoderm and advancing differentiation and maturation of endoderm derivatives. Enforced expression of *MIXL1* in the endoderm-inefficient hiPSCs enhanced the propensity of endoderm differentiation, suggesting that modulation of key drivers of lineage differentiation can re-wire hiPSC to the desired lineage propensity to generate the requisite stem cell products.

## Introduction

Human induced pluripotent stem cells (hiPSCs) are noted for their ability to differentiate into a multitude of cell and tissue types ^1–6^. Many protocols have been developed to direct the differentiation of hiPSCs to desirable types of endoderm cells or tissues, including the definitive endoderm^7–10^ and endoderm derivatives such as intestinal cells ^11–13^ pancreatic cells^14^ and hepatocytes^15–18^. A recent study of a bank of hiPSC lines derived from 125 individuals revealed that hiPSC lines respond differently when directed to differentiate to definitive endoderm^19^, suggesting that there is innate heterogeneity in the propensity for endoderm differentiation among hiPSC lines, which has been linked to specific quantitative trait loci (QTL). Both genetic determinants ^20–22^, and somatic or epigenetic memory related to the cell/tissue of origin^23–26^, have been shown to underpin the variable propensity of lineage specification and differentiation of hiPSCs. The impact of epigenetic memory on establishment of functional tissue nevertheless remains unresolved, even when examining cell lines from the same cellular resource and reprogrammed under the same conditions^27^.

In the present study, we investigated the propensity of endoderm differentiation of eleven lines of four sets of hiPSCs, each of which was derived from an independent cellular source, by tracking the differentiation from pluripotent cells to definitive endoderm (DE), hepatocytes and intestinal organoids (hIO). We showed that in these hiPSCs, early activation and a high level of *MIXL1* activity were associated with enhanced propensity of endoderm differentiation. In the mouse embryo, *Mix/1* is expressed in the primitive streak and the nascent mesoderm during gastrulation and expression persists in the primitive streak of the early-somite-stage embryo^28,29^. Loss of *Mix/1* function is associated with deficiency of DE and the local sequestration of the nascent mesoderm shortly after emergence at the primitive streak ^30^. In the mouse embryonic stem cells, loss of *Mix/1* function leads to inefficient differentiation of lateral mesoderm tissue and hematopoietic lineages^31^, whereas constitutive *Mix/1* activity promotes the differentiation of Foxa2+/ECad+ DE cells^32^. In mouse epiblast stem cells, activation of *Mix/1* at the early phase of differentiation correlates with efficient endoderm differentiation^28^. Analysis of the molecular attributes of DE differentiation revealed that the activity of *MIXL1* at the early phase of hiPSC differentiation promotes the differentiation of FOXA2+/SOX17+ DE cells in micropatterns^33^. We further showed that augmented expression of *MIXL1* in hiPSCs enhances endoderm propensity, offering new learning of how lineage propensity can be re-wired to generate fit-for-purpose pluripotent stem cells for translational application.

## Results

### Variable efficiency of definitive endoderm differentiation in hiPSCs

Eleven hiPSC lines from four cellular sources of different genetic backgrounds (two males and two females – Supplementary Table 1)^34,35^ were subjected to DE differentiation by following the STEMDiff definitive endoderm protocol (Figure 1A, B). To evaluate the developmental trajectory, cells were collected in triplicate cultures daily from Day 0 (pluripotency) to Day 4 (DE formation), and the expression of 88 genes involved in regulating pluripotency to germ layer differentiation was profiled (Supplementary Table 2). Line C32 showed the least progression across the 4 days of differentiation in the PCA plot (Figure 1C). To rank DE differentiation efficacy, we used the PC1 score as a proxy^19^, to infer an endoderm specification pseudotime. The average of triplicate PC eigenvalue along the PC1 axis was calculated to rank the hiPSC lines (Figure 1D). The results show that C32 and C7 ranked lowest while C9 and C11 ranked at the top for efficiency of endoderm differentiation. In addition, we observed that the cell lines are positioned more closely with members of the same group than with those of other three groups, suggesting a stronger effect of the genetic background of the cell lines on the efficiency of DE differentiation. In subsequent studies, we focused on cell lines of two groups: low endoderm propensity (C7 and C32) versus high endoderm propensity (C9, C11 and C16).

**Figure 1.**
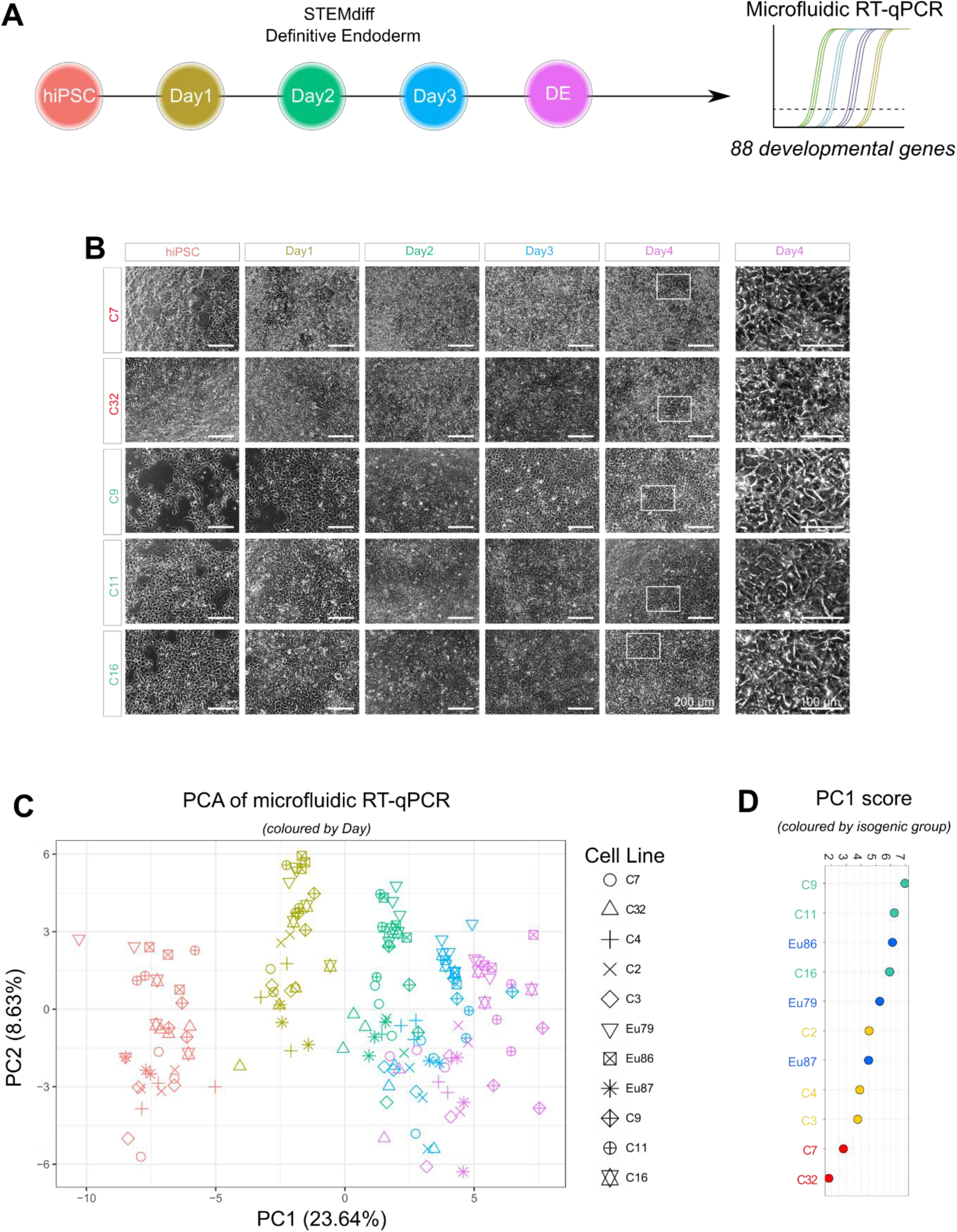
Endoderm differentiation of the four sets of isogenic lines: **A)** Protocol of definitive endoderm (DE) differentiation of hiPCS cells and profiling the expression of lineage-related genes by microfluidic-qPCR; **B)** Brightfield images of hiPSC cells at each day of the DE differentiation of 5 exemplar cell lines. Zoomed-in Day 4 image (right column) show that cell morphology is comparable among the cell lines. **C)** PCA of microfluidic RT-qPCR data of time-course DE differentiation. Each day is color-marked in the same scheme of panel A). **D)** PC1 score of eleven hiPSCs (colored by isogenic set) at Day 4 of DE differentiation presenting efficiency of differentiation as pseudotime.

These 5 cell lines were assessed for expression of FOXA2 and SOX17 on Day 4 of DE differentiation (Supplementary Figure 1A, B). Among the hiPSCs, C32, the lowest ranked based on the microfluidic RT-qPCR data (Figure 1D), displayed the lowest expression of both endoderm transcription factors (Supplementary Figure S1B), despite that the morphology of the C32 cells was comparable to other cell lines (Figure 1B), suggesting that C32 line is less efficient to differentiate into definitive endoderm.

### Low endoderm propensity line fails to generate advanced endoderm cell types

Hepatocyte (HCm) differentiation was used to further assess the endoderm propensity of C32 and C11 hiPSCs (Figure 2A). Both cell lines were able to differentiate into hepatocytes (Figure 2B and C). Microfluidic RT-qPCR analysis of genes specific to hepatocyte development (Supplementary Table 1) did not reveal any major differences in the transcriptome between these two cell lines at the hepatocyte stage of differentiation (Figure 2B, C). Although, differences can be spotted during DE differentiation where the C32 appears to be delayed in differentiation compared to C11. The phenotype of AAT- and ALB-expressing hepatocytes was also similar (Figure 2B, D; Supplementary Figure S1C). However, C32-derived hepatocytes showed lower Cytochrome P450 3A4 activity (Figure 2E) indicating that C32 hepatocytes might have different physiological attributes compared to C11 hepatocytes.

**Figure 2.**
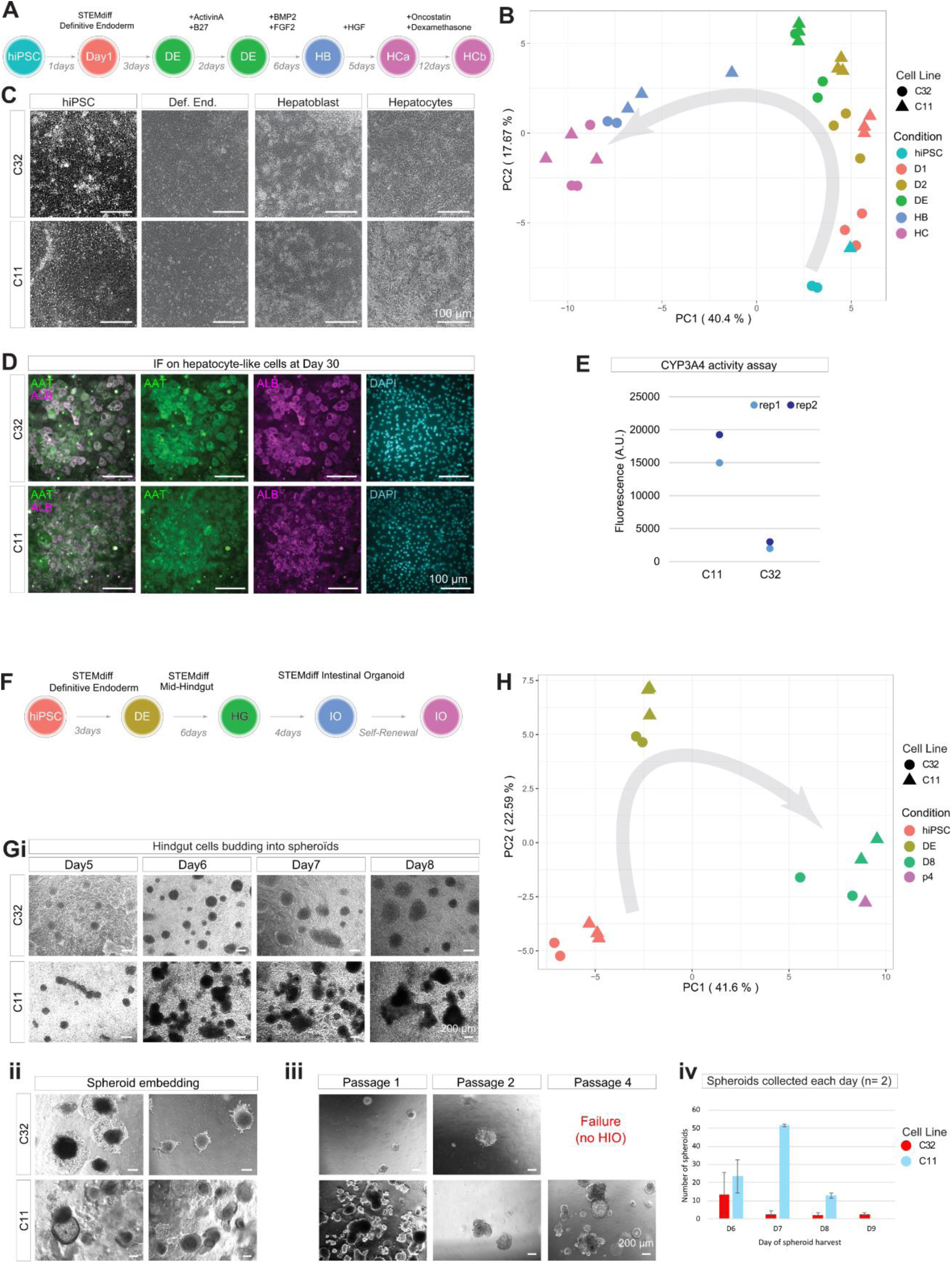
Low endodermal propensity cell line fails to produce functional tissue: **A)** Protocol for hepatocyte differentiation of C32 and C11 hiPSCs; **B)** PCA of microfluidic RT-qPCR data on hepatocyte differentiation, the grey arrow indicates pseudotime; **C)** Brightfield pictures of hepatocyte differentiation; **D)** Immunostaining images on AAT (green) and ALB (magenta) markers of hepatocyte differentiation; **E)** Fluorescence analysis of CYP3A4 activity; **F)** Protocol for generating intestinal organoids from C32 and C11 hiPSCs; **G)** Brightfield images of intestinal organoid development at (i) spheroid generation stage, (ii) the stage following embedding spheroids in Matrigel,(iii) intestinal organoids at passage 4 (iv) Quantification of budding spheroids collected for Matrigel embedding; **H)** PCA of microfluidic RT-qPCR data on intestinal organoid differentiation, the grey arrow indicates pseudotime; **I)** Immunostaining of intestinal organoids of C11 cell lines showing cell types of the gut epithelium: Goblet cells (UEA-1), Intestinal Stem cell (SOX9), enteroendocrine cells (CHGA), epithelium (CDX2) and Paneth cells (LYZ) and proliferating cells (Ki67). Nuclei stained by DAPI. Images on the left-most column showing the morphology of the organoid, images in columns 2 to 4 are zoomed-in of boxed region in column 1, showing the cell types.

The efficiency of producing human intestinal organoids (hIOs) was also examined in C32 and C11 lines (Figure 2F). At the budding spheroid stage, the C32 line generated fewer spheroids than the C11 line (Figure 2Gi, quantified in figure 2Giv). Microfluidic RT-qPCR analysis of the C32 hIOs did not show any discernible aberrant differentiation of cell types up to the spheroid stage (Figure 2H). Further growth of the C32 spheroids after embedding in Matrigel was impaired, compared to the robust growth of C11 spheroids (Figure 2Gii), and C32 spheroids did not progress beyond passage 3 (Figure 2Giii), pointing to inefficient generation of precursor cells at the budding spheroid stage for the intestinal organoids. In contrast, C11 derived hIOs continued to grow, accompanied by the establishment of typical cell types of the intestinal epithelium: enterocytes (CDX2+), intestinal stem cells (SOX9), enteroendocrine cells (CHGA), goblet cells (UEA-1) and Paneth cells (LYZ), and robust Ki67 expression, indicative of active proliferation and proper differentiation of the intestinal cells (Figure 2I). The C11 organoids could be grown, frozen and induced for further differentiation indicating that they are functionally competent. The hIO differentiation of C32 line was re-examined with changes in seeding densities and batches of cells, but the outcome remained poor.

Together, these results indicate that C32 line has an inherently low propensity for differentiation into definitive endoderm and advanced endoderm derivatives, and could not contribute to gut morphogenesis.

### Low propensity cell line displays a unique molecular signature during differentiation to primitive-streak like cells

We further investigated the low endoderm propensity C32 line and the high endoderm propensity C11 and C16 lines by single-cell RNAseq at 3 time points during differentiation: Day 0 (pluripotency), Day 1 (early germ layer) and Day 4 (definitive endoderm) (Figure 3A). After filtering, 41336 cells were retained for analysis. The tSNE plot (Figure 3B) showed that cells were segregated by time into three major clusters corresponding to each day. Interestingly, C32 cells displayed a unique transcriptomic profile at Day 1 and Day 4 (Figure 3C).

**Figure 3.**
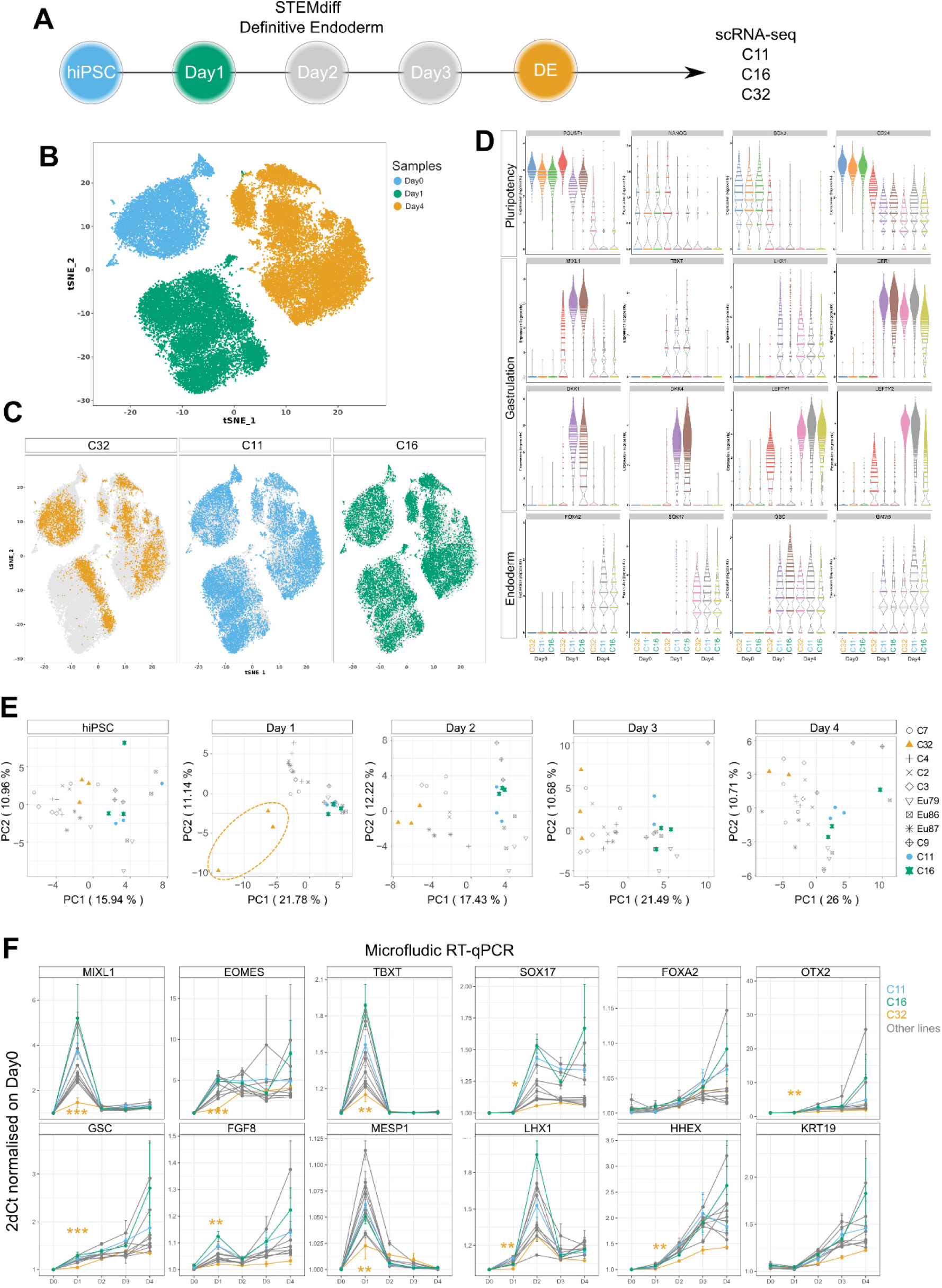
Single-cell transcriptome and gene expression profiles showing C32 is among the low propensity cell lines: **A)** Definitive Endoderm (DE) Differentiation protocol for C32, C16 and C11 lines. hiPSCs (blue), Day1 (green) and DE (orange) stages were collected for scRNAseq; **B, C)** tSNE plot of scRNA-seq data B) by day of *in vitro* differentiation and C) by individual cell lines; **D)** Expression pattern of genes associated with pluripotency, gastrulation and endoderm during *in vitro* DE differentiation; (**E**) PCA of microfluidic RT-qPCR data on DE differentiation, identical dataset as Figure 1D plotted for each day. **F)** Microfluidic RT-qPCR data of gene expression by time course during DE differentiation. C16, C11 and C32 are highlighted. p.value: significant differences between C32 versus at least C11 and C16 where * < 0.05, ** < 0.01, *** < 0.001.

To appreciate the pattern of cell clustering, the identity of single cells was annotated based on a human post-implantation embryo dataset^36^, as Epiblast, Primitive Streak, Nascent Mesoderm, Advanced mesoderm and Endoderm (Supplementary Figure S2A, B). Cells of the C32 line (55% of cells) retained an Epiblast signature at Day 1 and did not display a primitive-streak like cell state as the C11 and C16 cell lines (Supplementary Figure S2C, D). By Day 4, C32 cells have a signature reminiscent of the mesoderm.

Comparison of cell states markers’ expression for Days 0, 1 and 4 of DE differentiation (Figure 3D) showed C32 cells maintained high expression of pluripotency-related factors, *(SOX2, NANOG, POU5F1*) at Day 1. Genes associated with primitive streak differentiation, *MIXL1, LHX1, DKK1, DKK4*, and endoderm related genes (*GSC, GATA6*) were down-regulated at Day1 in C32 line. Interestingly, Nodal targets and antagonists, *LEFTY1* and *LEFTY2*, were up-regulated at Day 4, suggesting a downward modulation of the TGFβ pathway activity, which is necessary for DE differentiation.

To reinforce this inference, we further mined the microfluidic RT-qPCR data. Displaying the data of Figure 1D as a PCA for each day shows differences among cell lines that may not be visible on the full timeline of differentiation (Figure 3E). The C32 line displayed disparity on the main PC axis from the rest of the cohort at Day 1 (Figure 3E – Day1). The low differentiation efficiency does not appear to be linked to a slower down-regulation of pluripotency factors as there are no significant differences among the 11 cell lines for *POU5F1, NANOG, SOX2* and *ZFP42* mRNA expression kinetics (Supplementary Figure S3Ai). *TDGF1*, a gene coding for NODAL co-receptor, is significatively down-regulated and FGF5 is overexpressed in C32 compared to C11 and C16 at Day 1.

Gastrulation genes such as *MIXL1, GSC, TBXT, EOMES, MESP1* and *FGF8* are all down-regulated at Day 1 in C32 compared to the cohort (Figure 3F). C32 also has the lowest levels of expression of *SOX17, OTX2, GATA4, AFP* and *AXIN2* at Day 4 (Figure 3F and S3Aii). Surprisingly, *FOXA2* and *NODAL* are not dysregulated in the C32 line. In addition, mesodermal genes (*BMP4, MYH7, KLF5* and *CD34*) (Supplementary Figure S3Aiii) as well as ectodermal genes (*KRT10, SOX1, NES, FOXD3, PAX6* and *DCX*) (Supplementary Figure S3Aiv) showed no significant difference between C32 and other lines. *VIM*, a gene coding for cell adhesion protein, is significantly more expressed in C32 (Supplementary Figure S3Av). Finally, we could see *PDGFRa* and *KDR (FLK1)*, two targets of *MIXL1* in mouse ESC^37^, down-regulated in C32 at Day 1 of differentiation (Figure 3F and S3Aii).

The lowest expression of *SOX17* at Day 4 (Figure 3F) and the reduced proportion of endoderm cells in the C32 line relative to C11 and C16 cell lines (Figure S2D) is in agreement with the immunofluorescent data (Figure S1B) and the low efficiency in producing mature tissue (Figure 2).

### Hippo signaling and cell adhesion in the low efficiency cell line

To mine the transcriptomic differences at Day 1 if DE differentiation, bulk RNA-seq analysis was performed and has revealed differential regulation of gene activity between the low (C7, C32) and the high propensity lines (C9, C11, C16). Transcriptomic differences showed that C32 differs significantly from the other cell lines (Supplementary Figure S3B, C, D), displaying enrichment of transcriptomic signature of mesoderm (circulatory system and heart: *RUNX1, VEGFA*) and ectoderm (neural: *SOX2, POU3F2*) (Supplementary Figure S3C) derivative. Genes associated with gastrulation (e.g.; *MIXL1, EOMES, MESP1, APLN, DKK1, GATA6*) (Supplementary Figure S3C) were down-regulated in C32. C7, a hiPSC line of the same group as C32, which is more efficient in definitive endoderm differentiation (Figure S1B) expressed genes associated with gastrulation (*MIXL1, EOMES, TBXT, SNAI1, CER1*) at a higher level, whereas C32 showed enhanced expression of genes associated with cell adhesion (*VIM, EZR, FLNA, FLNC, COL1A1, FN1, ITGA2/3/6*) and Hippo signaling (*CCN1, CCN2, AMOT, AJUBA, CDH11*) (Supplementary Figure S3E).

Results of transcriptomic analysis highlight a bias in the lineage propensity for ectoderm and mesoderm and a low propensity for endoderm in germ layer differentiation of C32 line. These results further point to the potential involvement of Hippo signaling and cell adhesion in underpinning the different propensity of endoderm differentiation between cell lines of isogenic background such as C32 and C7.

Collectively, scRNA-seq, bulk RNA-seq and microfluidic RT-qPCR all pointed at a delay or inefficient differentiation of the C32 cells to primitive-streak like cells.

### Single-cell transcriptomic analysis as a tool to rank lineage propensity

To test if the findings from this cohort of iPSCs may be extrapolated to other hiPSC lines, the scRNAseq dataset was combined with that of 125 hiPSC lines that have been profiled for endoderm differentiation *in vitro*^19^ (Figure 4A and B). Of note, this external dataset has been generated using a different library and sequencing technology (SMART-seq2) and differentiation protocol (homemade versus our study using the STEMdiff kit). Each cell line was ranked for endoderm propensity based on the PC1 eigenvalue at Day 4 (Figure 4C) similarly to Figure 1D. The data show that C32 ranks among the cell lines in the lowest 20% propensity, while C16 and C11 rank higher

**Figure 4.**
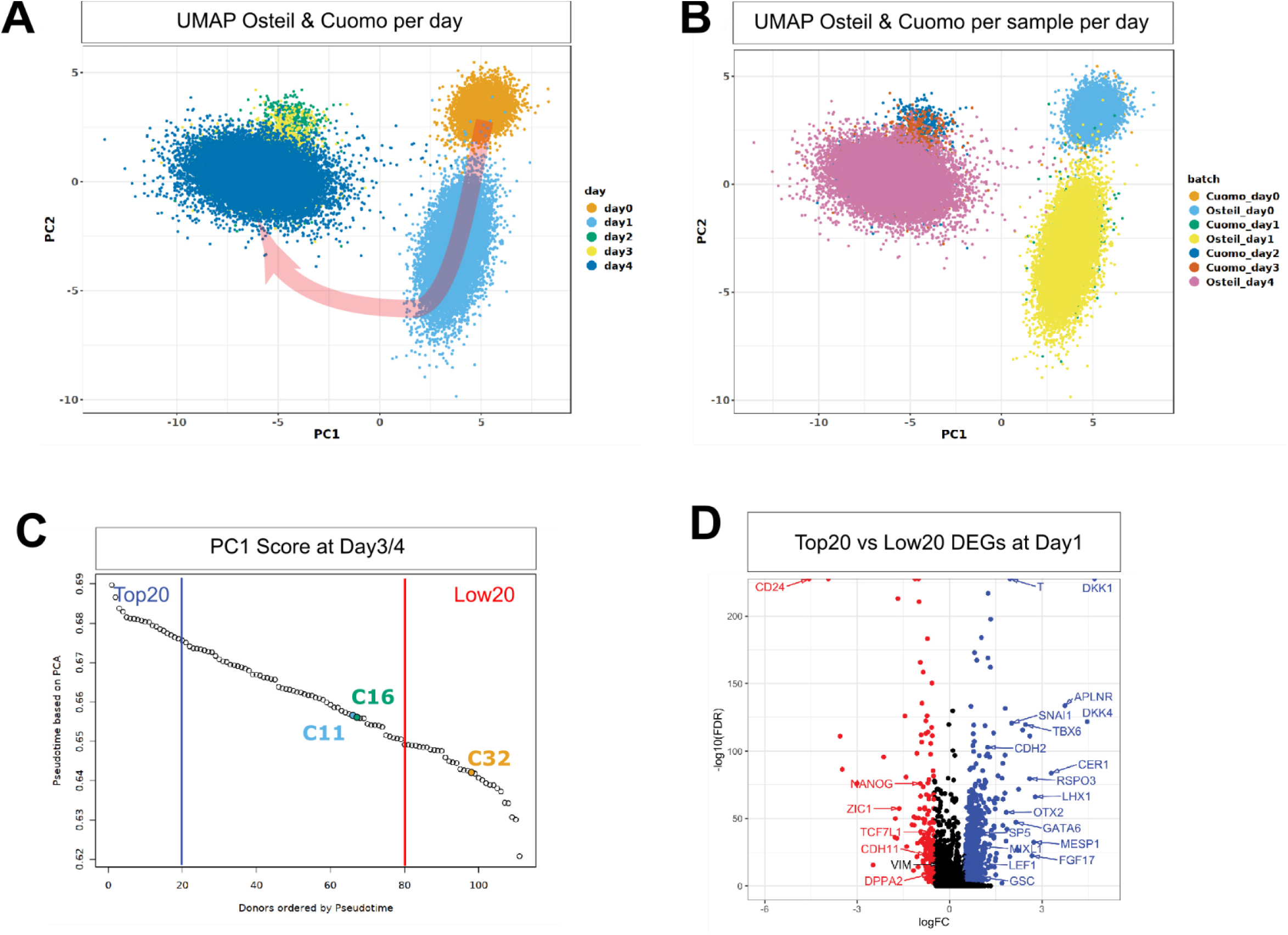
Ranking of iPSCs for endoderm propensity. **A and B)** PCA showing integration of our scRNA-seq data with data of Cuomo et al., **A)** represents cells grouped by day of *in vitro* differentiation, arrow indicates time course of differentiation; **B)** grouped by sample.; **C)** PC1 axis projection of combined hiPSCs from Cuomo et al. and current study at Day 3/4 of DE differentiation representing efficiency of differentiation as pseudotime. Position of C32, C11 and C16 cell lines from current study are highlighted; **D)** Differentially Expressed Genes (DEGs) between the Top 20 cell lines (more efficient) (blue) versus the Low 20 cell lines (less efficient) (red) at Day 1 of differentiation presented in volcano plot.

Using this classification, the highest 20 cell lines (Top20) and the lowest 20 cell lines (Low20), were analyzed for DEGs at Day 1 of differentiation. This analysis revealed that genes associated with primitive streak formation were up-regulated in Top20 cell lines (*TBXT, DKK1, SNAI1, MIXL1, LHX1, etc…*) while Low20 cell lines retained a pluripotent signature (*NANOG, DPPA2, ZIC1*) (Figure 4D). The data infer that the Low20 lines are less competent for primitive streak differentiation and that the C32 characteristics are shared by other hiPSCs of low propensity of endoderm differentiation.

In conclusion, irrespective of the sequencing technology used and the differentiation protocols, single cell RNAseq data can be used to rank hiPSC lines in the context of the efficiency of endoderm differentiation by PC1 eigenvalue score. This will enable the selection of cell lines with optimal differentiation propensity prior to using these cells to generate the endoderm derivatives for research and application.

### MIXL1 is required for promoting primitive streak differentiation

From the above analysis, it was found that the gastrulation transcriptomic program is not activated properly in those cell lines that are less efficient in differentiating into definitive endoderm. One gene in particular caught our attention, MIXL1, due to its role in endoderm formation in mouse embryos. To test if dysregulation of *MIXL1* expression early in lineage differentiation may underpin the low endoderm propensity, we next studied the requirement of *MIXL1* for endoderm differentiation in human stem cells. To create a loss-of-function setting, frameshift mutations of *MIXL1* were engineered in C32 and C16 lines by CRISPR editing to generate *MIXL1* loss of function cell lines: C32-MKO and C16-MKO. We verified the *MIXL1* gene editing event by sequencing the targeted regions, and confirmed that the editing did not impact on the pluripotency of these lines (Supplementary Figure S4A). To create a gain-of-function setting, C32 cell line was engineered to constitutively express the dead Cas9, with no nuclease activity, linked to VP64, a potent transcriptional activator (dCas9-PVP64)^38,39^. This line was genetically modified further to express two sgRNAs that targeted the dCas9-VP64 to the promoter of *MIXL1*, in a doxycycline controllable fashion (C32-iMIXL1)^40^. Among the sqRNAs guides, sgRNA4 and 7 (Figure S4B), elicit the strongest activation of *MIXL1* in HEK cells (Figure S4C).

We next quantified *MIXL1* protein expression by immunofluorescence in the KO lines and the C32-iMIXL1 line with and without induction at Day 1 of DE differentiation (Figure 5A). No *MIXL1* expression can be detected in C32-MKO and C16-MKO (Figure B) relative to C32-iMIXL1 cells treated with DMSO (C32-iMIXL1:Dox0) (Figure 5B). Maximal induction of *MIXL1* expression was achieved, at a saturating concentration of 2*μ*g/mL Dox (C32-iMIXL1:Dox2) (Figure 5C). Beyond which, at 16*μ*g/mL Dox, *MIXL1* expression was reduced possibly due to the toxicity that impacts on mitochondrial gene activity. The levels of induced expression of MIXL1 with 1*μ*g/mL of Doxycycline (C32-iMIXL1:Dox1), quantified by immunofluorescence, were within the physiological range of the high propensity C16 line at Day 1. Although we noticed a higher level of MIXL1 expression in C32-iMIXL1:Dox0 compared to the founder line, C32, indicating potential leakiness of the construct, albeit the level remains below that of C16.

**Figure 5.**
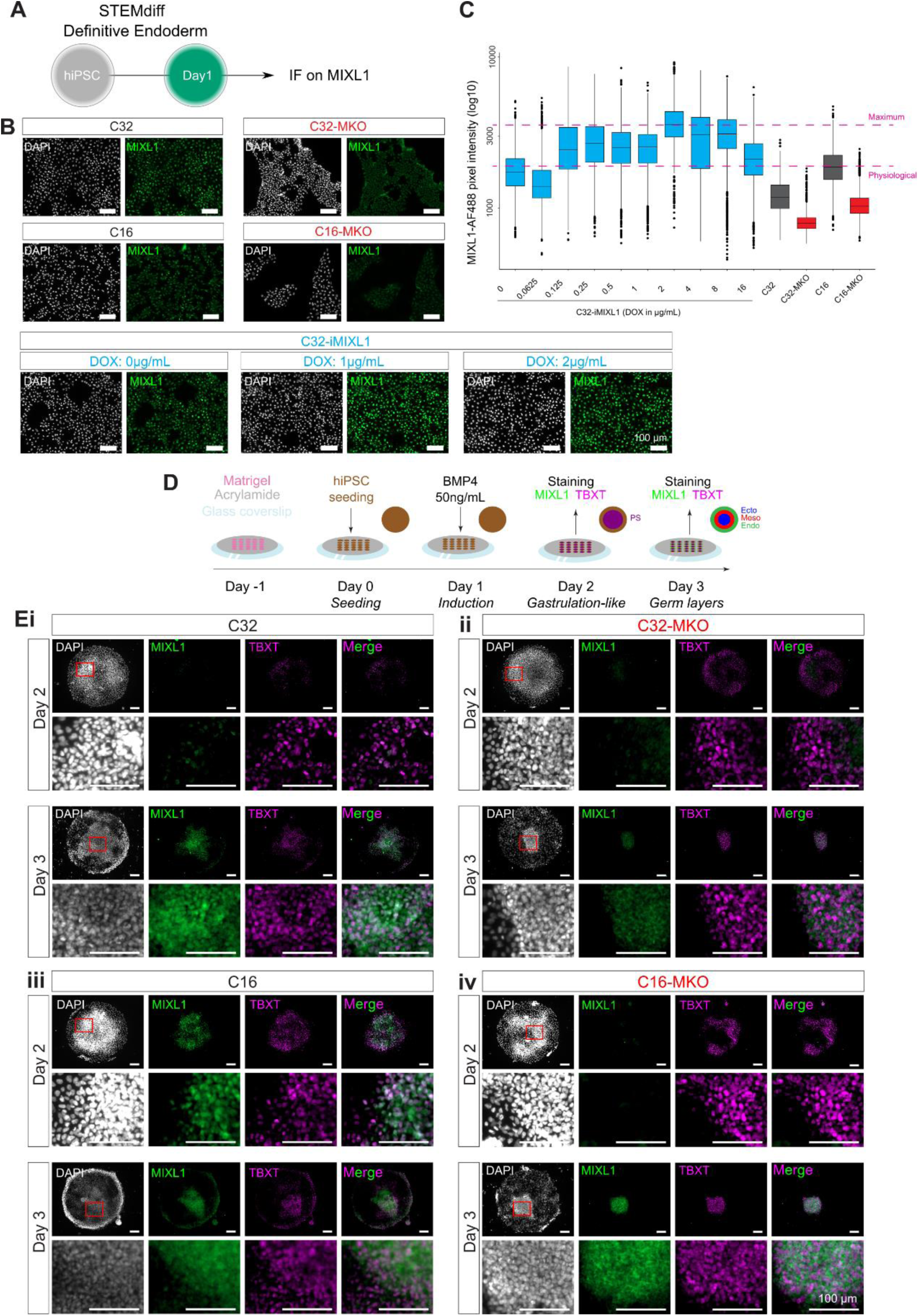
MIXL1 functional genomic study: **A)** Experimental workflow where definitive endoderm differentiation protocol was stopped at Day 1 and cells stained for MIXL1 via immunofluorescence (IF); **B)** Immunostaining images on MIXL1 (green) for C16 and C32 either WT or MIXL1-KO (MKO) and C32 with inducible MIXL1 expression (C32-iMIXL1) under three different concentrations of doxycycline (0, 1 and 2µg/mL). Nuclei are revealed by DAPI (white). **C)** MIXL1 signal intensity normalized on DAPI signal for concentration of doxycycline spanning from 0 to 16µg/mL, WT and MKO conditions. 2 levels are highlighted: a physiological level, corresponding to the level of C16 and 1µg/mL of doxycycline and an overexpression level corresponding to 2µg/ml. (n= 3); **D)** Protocol for stem cell micropattern model of germ layer differentiation where MIXL1 and TBXT were stained 24h (Day 2) and 48h (Day 3) after BMP4 treatment; **E)** Immunostaining images of MIXL1 and TBXT at 24h and 48h after BMP4 induction on C32 **(i)** C32-MKO **(ii)**, C16 **(iii)**, and C16-MKO; **(iv)** Zoomed-in regions from red highlighted square are shown in the row underneath.

To assess the function of *MIXL1*, we used the 2D stem cell micropattern model to elucidate its functional attribute in germ layer differentiation. Four cell lines, C32, C32-MKO, C16 and C16-MKO, were used to generate micropatterned cultures that recapitulate germ layer formation in response to BMP4^33,41,42^. The emergence of primitive streak-like cells was assessed via the immunostaining of TBXT and MIXL1 proteins 24h and 48h after BMP4 supplementation (Figure 5D). In the C32 cell line at 24h MIXL1 signal was almost undetectable and only increased at 48h (Figure 5Ei). In the C32-MKO line, MIXL1 was not detected at 24 h and was at background level in the cytoplasm at 48h (Figure 5Eii). TBXT expression stayed at basal level in both C32 and C32-MKO lines (Figure 5Ei, ii). In the C16 line TBXT and MIXL1 were expressed at peak levels in the nuclei at 24h (Figure 5Eiii) followed by decreased expression at 48h (Figure 5Eiv), with some cells co-expressing both TBXT and MIXL1. In C16-MKO line, MIXL1 is detected as background in the cytoplasm (Figure 5Eiv).

These data suggest that in a micropattern model, C32 is delayed in activating MIXL1. Moreover, TBXT pattern of expression is conserved in C32 and MKO conditions.

### MIXL1 plays a role in chromatin organization

To elucidate the impact of MIXL1 on chromatin accessibility, Assay for Transposase-Accessible Chromatin using sequencing (ATAC-seq) was performed on C16 and C16-MKO cell lines at Day 1 of DE differentiation, when *MIXL1* expression is maximal, in an DE efficient cell line. Comparing the accessible regions (or reads pile-up called as peaks) showed that *MIXL1* deletion led to multiple changes in chromatin accessibility (Supplementary Figure S5A), and in particular, less closing regions (Supplementary Figure S5B), than opening (Supplementary Figure S6C). This observation suggests that MIXL1 may be directly or indirectly responsible for opening and closing regions during germ layer differentiation.

To understand the role of these regions, motif discovery was performed on differentially accessible chromatin regions. This analysis revealed that peaks with less accessibility in MKO lines contain the motifs TAATNNNATTA (PROP1, PHOXA2), which is the dual homeobox motif recognized by MIXL1 (Supplementary Figure S5D). In the absence of *MIXL1*, more accessible peaks are associated with TEAD1 and FOXH1 motifs (Supplementary Figure S5E). FOXH1 is known as a cofactor of GSC which negatively regulates MIXL1 in the mouse^43^. TEAD1 is the transcription factor bound by YAP/TAZ when Hippo signaling is inhibited^44^ suggesting MIXL1 activation may lead to close TEAD1 accessible region, finding that is consistent with the observation that the C32 cell line which expresses low *MIXL1* exhibits up-regulation of YAP/TAZ targets (*CCN1*, *CCN2*) (Supplementary Figure S3E).

Collectively the data reveal that MIXL1 transcription factor may be involved in modulating chromatin accessibility of its targets and possibly influences signaling pathways such as Hippo and WNT, as inferred from the TCF3 motifs found in open regions in the C16-MKO lines.

### Physiological levels of MIXL1 activity can enhance the endoderm propensity of low efficiency iPSCs

To assess how outcomes of endoderm differentiation are modulated by different levels of *MIXL1* expression, FOXA2 and SOX17 expression was quantified after 4 days of differentiation in MKO and iMIXL1 lines (Figure 6A). Increasing MIXL1 expression in C32-iMixl1 resulted in higher expression of both markers (Figure 6B, C), further reinforcing a role of *MIXL1* in promoting endoderm differentiation. Surprisingly, however, C32-MKO cells displayed FOXA2 and SOX17 expression levels similar to C16-MKO and C32-iMIXL1:Dox0. This finding suggests that MIXL1 dysregulation may not be the sole cause of inefficient endoderm specification and differentiation. Another explanation may be the cloning step of C32-MKO involving a single cell regenerating an entire cell line. This step may have led to the loss of the properties of the founder C32 line. However, these results indicate that modulation of MIXL1 expression can have an effect on DE formation.

**Figure 6.**
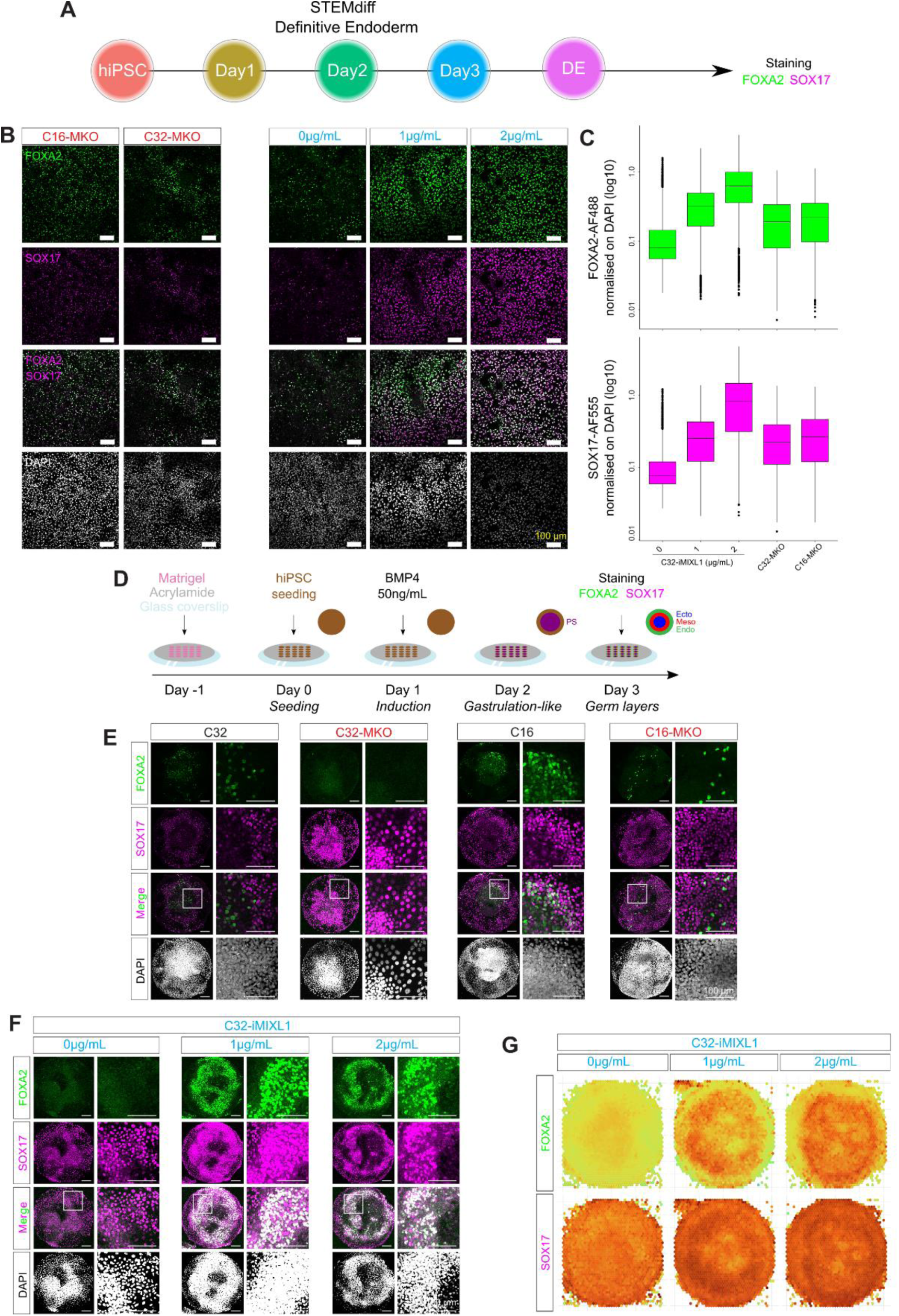
MIXL1 induction rescues DE phenotype. **A)** Protocol of definitive endoderm differentiation and analysis of marker gene expression at Day 4 of differentiation cells; **B)** Immunofluorescence detection of FOXA2 (green) and SOX17 (magenta) on Day 4 of DE differentiation in C16-MKO, C32-MKO and C32-iMIXL1 cell lines at three different concentration of doxycycline; **C)** Quantification of FOXA2 and SOX17 immunofluorescence signals from images on panel B; **D)** Protocol for micropattern modelling of germ layer differentiation; **E, F)** Immunofluorescence detection of FOXA2 (green) and SOX17 (magenta) in E) C32, C32-MKO, C16 and C16-MKO lines by 48h (Day3) after BMP4 induction, and **F)** C32-iMIXL1 under three different concentrations of doxycycline (0, 1 and 2µg/mL by 48h after BMP4 induction; **G)** Display of the increasing signal intensity, measured digitally, for endoderm markers in C32-iMIXL1 micropatterns induced by doxycycline at 0, 1 and 2µg/mL (n= 10 for each dosage).

DE differentiation was also assessed by FOXA2 and SOX17 expression in micropatterns, by 48h after BMP4 induction (Day 3 - Figure 6D). FOXA2+/SOX17+ cells were organized as an inner ring in the C16 line (Figure 6E). However, no double positive cells were detected in C32 line (Figure 6E). Instead, separate populations of FOXA2+ cells (inner domain) and SOX17+ cells (outer domain) were identified (Figure 6E).

These results show that a change in culture format did not alter the outcome of DE differentiation in the low efficiency C32 line still showing defects in forming FOXA2+/SOX17+ cells. In MKO lines, the loss of MIXL1 activity resulted in a reduced population of FOXA2+ cells while SOX17 expression did not change (Figure 6E, Supplementary Figure S6).

To elucidate the effect of induced *MIXL1* activity on DE differentiation of the low propensity iPSC line, the C32-iMIXL1 lines were cultured in micropatterns under BMP4 condition with induction of physiological level of *MIXL1* activation (DOX1: 1µg/mL) and with *MIXL1* over-expression (DOX2: 2µg/mL). Endoderm differentiation at Day 2 of differentiation of induced C32-iMIXL1, compared with C32 line (parental low propensity line) and C16 line (high propensity line), was assessed by the presence of SOX17+ and FOXA2+ cells in the micropatterns. C32-iMIXL1 cells displayed increased number and density of SOX17+ and FOXA2+ cells with DOX induction, compared to C32 line and C16 line and the control condition without doxycycline (Figure 6E - G). Inducing MIXL1 above physiological level (C32-iMIXL1:Dox2) only slightly increase the population of double positive cells (Supplementary Figure S7).

Reconstitution of physiological levels of MIXL1 activity in the inherently inefficient C32 line therefore enhanced the endoderm propensity of iPSCs. In addition, increasing the doxycycline concentration raised the number of double positive cells in the micropatterned cultures, indicating a causal effect between *MIXL1* expression level and the efficiency of definitive endoderm differentiation.

## Supporting information

Cell line meatadata

Microfluidic oligos

ATACseq oligos

## Acknowledgments

We acknowledge the support of CMRI Advanced Imaging Facility for all imaging and imaging analysis and CMRI Vector and Genome Engineering Facility for generating the KO lines, Prof. Kristopher Kilian and his team for sharing the micropattern protocols and training. This work was supported by the National Health and Medical Research Council (NHMRC) Project Grant (ID1127976). PPLT was supported by the NHMRC Senior Principal Research Fellowship (Fellowship Grant ID110751)

## Author contributions

PO, EW and PPLT conceived of the study. PO, SW, Ni.S, PPLT designed experimentation. PO and Ni.S conducted the cell and molecular biology, imaging, and biochemical analysis with assistance by AQ, Jy.S and XBL. SW conducted the hepatocyte experiments and the engineering of the C32-iMIXL1 cell line. HK and Ni.S conducted the RNA-seq and scRNA-seq and NA performed the bioinformatic analysis. SC assisted with preliminary data analysis. Ni.S conducted ATAC-seq and Na.S the data analysis. PO conducted experimental analysis and data interpretation and prepared the data for publication with assistance by NA and PPLT. PO, Ni.S and PPLT wrote the manuscript with input from all authors.

## Discussion

In this study, human induced pluripotent stem cell (hiPSC) lines were analyzed and compared for their propensity in generating definitive endoderm that is capable of progressing to functional endoderm derivatives, here tested by the formation of intestinal organoids and hepatocytes. One cell line of the cohort (C32) was found to be inefficient in differentiation towards the endoderm lineage. The low propensity for endoderm derivatives is accompanied by a bias toward the mesoderm lineage. Our results suggest that during germ layer differentiation, C32 activates the genetic program of vascularization and heart formation more efficiently than the rest of the cohort (See Supplementary Figure S3C). This is corelated to the mouse MIXL1-/- phenotype exhibiting increase mesodermal production and no endoderm.^30^ In addition, the C32 hiPSC line has been used in the generation of kidney organoid^6^, containing predominantly mesodermal derivatives, thereby consistent with mesoderm bias of the C32 cell line.

We sought to determine molecular markers of endoderm propensity of hiPSC lines, at the pluripotency and early exit stages. Transcriptomic analyses did not reveal evident bias in lineage propensity of the inefficient C32 line at these stages. The retained pluripotency level may underpin the poor performance in endoderm differentiation of this cell line. To gain a holistic view of the differentiation potential of hiPSC lines, 4 groups of hiPSC lines (2 males and 2 females) were subjected to transcriptomic analysis. This further highlighted that in C32 line, endoderm differentiation is inefficient which is accompanied by delay in the initiation of germ layer differentiation, compared to other cell lines in the cohort, as well as to the more efficient cell line of the same isogenic group.

The single cell transcriptomic data was analyzed in conjunction with the transcriptomic data of a large-scale study^19^ surveying 125 hiPSC lines during definitive endoderm differentiation. Despite the use of different protocols and sequencing chemistry, the findings were remarkably comparable. Cell lines with the lowest differentiation score at the end of the DE differentiation, have a significantly low expression of genes involved in gastrulation including *MIXL1*. These results further validated our microfluidic RT-qPCR findings and enabled the identification of a gene panel and development of a novel tool for ranking hiPSC lines by the propensity of endoderm differentiation and ability to generate mature endoderm tissues.

The transcriptomic survey of the cohort revealed that the gene *MIXL1* is expressed at a low level in endoderm-incompetent cell lines, and this is corroborated via cross comparison of the aforementioned study^19^ (Figure 4D). Our endoderm differentiation data of stem cell-derived micropattern further points to a causal relationship between the expression of *MIXL1* and efficiency of DE differentiation. We proposed that *MIXL1* is a useful molecular marker, through quantification of expression in cells at 24h of directed differentiation, for screening human pluripotent stem cells for competency of endoderm differentiation.

The ATAC-seq data pointed to a possible role for MIXL1 in regulating chromatin accessibility which was not previously noted. The target regions opened in the MIXL1-KO cell line are strongly enriched for TEAD1 and TCF3 binding sites. TEAD1 is a transcription factor involved in Hippo signaling^44^. This data, taken together with the finding that Hippo target genes (e.g., *CCN1, CCN2*) are activated in the inefficient C32 line, suggests a relationship between chromatin status and the Hippo pathway that is regulated by *MIXL1*. On the other hand, TCF3 containing sites, a transcription factor bound by β-Catenin upon Wnt activation, appear to be more accessible in MIXL1-KO condition. This indicates that in C32 line Wnt pathway is probably differentially activated, which may account for the enhanced differentiation of mesodermal derivatives^45^ Interestingly, the closed chromatin region in the MIXL1-KO line mainly contains the dual homeobox binding sites TAATNNNATTA, recognized by MIXL1. This suggests that MIXL1 may be involved in regulating accessibility to its own transcriptional targets.

Finally, our study demonstrated a causal link between *MIXL1* activation and definitive endoderm differentiation in human pluripotent stem cells. Mouse studies have shown that MIXL1 overexpression led to increase mesoderm and endoderm^32,46^, but these studies were using constitutive overexpression of *Mix/1*. Here we demonstrate that temporal and dose control of the expression of *MIXL1* is instrumental to the differentiation of definitive endoderm. Our study has provided learnings of the regulation of endoderm differentiation in a cohort of hiPSCs, establishing a framework for future study of the molecular mechanisms that underpin endoderm specification during germ layer differentiation of pluripotent stem cells and in human embryonic development.

## Resource availability

### Lead contacts

Further information and requests for resources and reagents should be directed to and will be fulfilled by the lead contacts, Pierre Osteil or Patrick Tam.

### Material availability

The materials used in this study are listed in the key resources table. Materials generated by our laboratory in this study are available on request, however, there are restrictions to the availability of human iPSC lines due to a Material Transfer Agreement.

### Data and Code availability

- All raw sequencing data can be found on Gene Expression Omnibus under the accession number GSE260552 (ATAC-seq), GSE260553 (RNA-seq), GSE260554 (scRNA-seq) and are available as of the date of publication.
- All microfluidic RT-qPCR data can be found on the GitHub page (https://github.com/PierreOsteil/ScriptsForOsteilEtAl2024) and are available as of the date of publication.
- All original codes are available as of the date of publication and can be found on the following GitHub page: https://github.com/PierreOsteil/ScriptsForOsteilEtAl2024. Bioinformatic source codes and their corresponding DOIs are listed in the key resources table
- Any additional information required to reanalyze the data reported in this paper is available from the lead contact upon request.

## Experimental Model and Subject Details

### Cell Lines

A cohort of 11 human iPSC lines composed of 2 to 3 isogenic cell lines from 4 patients (2 males and 2 females) was provided by the Australian Institute for Bioengineering and Nanotechnology (AIBN), University of Queensland. Briefly, hiPSC lines derived from fibroblast cells or foreskin tissue were generated using a non-integrating episomal reprogramming system (oriP/EBNA1-based pCEP4 episomal vectors pE-P4EO2SCK2MEN2L and pEP4EO2SET2K from Addgene) carrying *OCT4, SOX2, KLF4* and *CMYC*. All lines maintained a normal karyotype and were capable of forming teratomas that contained derivatives of the three germ layers^34,35^ (Summarized in Supplementary Table 1). For routine maintenance, hiPSCs were cultured in mTeSR1 (StemCell Technologies) on six-well plates precoated with hESC-qualified Matrigel (Corning). The culture plates were incubated at 37°C and 5% CO_2_. The medium was changed daily. The colonies’ morphology was evaluated under an inverted microscope. Cells were passaged at 70-80% confluency with ReLeSR (StemCell Technologies) at a split ratio of 1/5 to 1/30 depending on the cell line, into a new well of a 6-well plate. Experiments were approved by the Sydney Children’s Hospitals Network Human Research Ethics Committee under the reference: HREC/17/SCHN/167.

## Method details

### Differentiation protocols

#### Definitive endoderm differentiation and characterization

For direct differentiation into Definitive Endoderm, the cells were subject to induction using STEMdiff Definitive Endoderm kit (StemCell Technologies) for 4 days, following manufacturer’s protocol. Briefly, cells were passaged into single cells with StemPro Accutase Cell Dissociation Reagent (Life Technologies) and seeded with mTeSR1 containing Y-27632 dihydrochloride Rock Inhibitor (Tocris). After 24 h, the cells are washed with Dulbecco’s phosphate buffered saline (dPBS) (Gibco) and then cultured for 4 days in STEMdiff Definitive Endoderm Basal medium with Supplements A and B for the first day and then Supplement B only for the subsequent 3 days, with daily medium changes. Samples were harvested daily for RNA and Protein extraction (n=3).

To characterise the definitive endoderm, cells were seeded onto glass coverslips coated with hESC-qualified Matrigel (Corning) before treating with STEMdiff Definitive Endoderm kit (StemCell Technologies) as described above. Cells were fixed in 4% paraformaldehyde in PBS at RT for 20 min. They were washed with PBS twice and then permeabilized with 0.1% Triton X-100 (Sigma-Aldrich) in dPBS (Gibco) (PBST) at RT for 5 min. The cells were blocked with 3% bovine serum albumin (Sigma-Aldrich) in PBST at room temperature for 1 h. They were incubated with primary antibody at 4°C overnight (SOX2 (Cell Signaling) 1:400, OCT4 (Santa Cruz) 1:50, FOXA2 (Abcam) 1:300, SOX17 (R&D Systems) 1:20). Cells were washed with dPBS three times, then incubated with corresponding secondary antibodies at RT for 1 h. The cell nuclei were stained with DAPI (1*μ*g/ml) (Thermo Fisher Scientific) in dPBS for 10 min at RT, and then washed three times with dPBS. Cells were mounted with Fluoromount-G (Thermo Fisher Scientific) and imaged on Ziess Axio Imager Z2 widefield microscope.

#### Human Intestinal Organoid differentiation and characterization

hiPSC-derived intestinal organoids were formed using the STEMdiff Intestinal Organoid Kit (StemCell Technologies), following the manufacturer’s protocol. Briefly, cells were passaged as clumps using ReLeSR (StemCell Technologies). Once 80-90% confluency was reached, differentiation was initiated with DE Medium (STEMdiff Endoderm Basal Medium plus STEMdiff Definitive Endoderm Supplement CJ) for 3 days, with daily medium changes. Subsequent mid-hindgut differentiation was induced with MH Medium (STEMdiff Endoderm Basal Medium plus STEMdiff Gastrointestinal Supplement PK and STEMdiff Gastrointestinal Supplement UB) for 6 days, with daily medium changes. Free-floating mid-hindgut spheroids, collected at 24 h intervals within the MH Medium treatment, were embedded in Basement Membrane Matrigel (Corning) in wells of NunclonDelta surface 24-well plate (Thermo Fisher Scientific), overlaid with STEMdiff Intestinal Organoid Growth Medium (STEMdiff Intestinal Organoid Basal Medium plus STEMdiff Intestinal Organoid Supplement (StemCell Technologies) and GlutaMAX (Gibco)), performing medium change every 3 - 4 days, incubating at 37°C with 5% CO_2_. After 7 - 10 days of incubation, cultures were passaged. Briefly, all plasticware were pre-wetted with Anti-Adherence Rinsing Solution (StemCell Technologies). Matrigel domes containing organoids were broken manually by pipetting up and down with cold DMEM/F-12 (Gibco), seeding 40-80 organoid fragments per 50*μ*l Matrigel dome.

To harvest organoids for immunofluorescence, they were removed from Matrigel similar to that described for splitting organoids above and were fixed in 4% paraformaldehyde in PBS at RT for 30 min. They were washed with PBS twice and then permeabilised with 0.1% Triton X-100 (Sigma-Aldrich) in dPBS (Gibco) (PBST) at RT for 1 h. The organoids were blocked with CAS-Block (Thermo Fisher Scientific) for 90 min and then incubated with primary antibody (Sox9 (Sigma-Aldrich) 1:500, Ki67 (Abcam) 1:250, CHGA (Novus Biologicals) 1:200, CDX2 (Biogenex) 1:250, Lysozyme (Dako) 1:200) overnight at 4°C. Organoids were washed with PBST four times, then incubated with corresponding secondary antibodies and stain (DAPI (1 *μ*g/ml) (Thermo Fisher Scientific) for nuclei in all samples and UEA-1 (Vector Laboratories) 1:200, for select organoids) in CAS-Block at RT for 3 h. The organoids were then washed four times with PBST, followed by clearing in FUNGI solution (50% (v/v) glycerol, 9.4% (v/v) dH_2_O, 10.6M Tris base, 1.1mM EDTA, 2.5M fructose and 2.5M urea) for 40 min. Organoids were imaged using a *μ*-slide (Ibidi) on Zeiss Cell Observer Spinning Disc confocal microscope.

#### Hepatocyte differentiation and characterization

hiPSC lines were differentiated toward hepatocytes following the protocol from Baxter et al^47^, with modification during the definitive endoderm stage. Briefly, hiPSC lines were plated onto hESC qualified Matrigel coated tissue culture plates at 131,000 cells/ cm^2^ (Day 0). After 24 hours, cells were washed with RPMI medium and differentiation protocol commenced. Briefly, on Days 1 - 4 cells were treated with STEMdiff Definitive Endoderm kit (STEMCELL Technologies) following manufacturer’s guidelines, to induce definitive endoderm (DE), followed by 2 days of treatment with RPMI media containing 100 ng/ml Activin-A (Peprotech) and 1:50 B-27 supplement minus insulin (Thermo Fisher Scientific), fed daily. Hepatoblast stage (HB) was 6 days (days 7-12) with 20 ng/ml BMP2 (Peprotech) and 30 ng/ml FGF basic (R&D Systems) in Hepatocyte Culture Medium BulletKit (HCM) (Lonza), fed daily. Hepatocyte induction (HCa) commenced (Days 13 - 17) with 20 ng/ml Hepatocyte Growth Factor (HGF) (Peprotech) in HCM (Lonza), fed every other day. Finally, Hepatocyte maturation (HCb) concluded with 10 ng/ml Oncostatin M (R&D Systems) and 100 nM Dexamethasone (Sigma-Aldrich), fed every other day for Days 18 - 29.

Hepatocytes (HCb) characterized by immunofluorescence were plated on Nunc Lab-Tek Chamber slides (Thermo Fisher Scientific) before differentiating. Cells were fixed in paraformaldehyde in PBS at room temperature for 10 min. The remaining protocol is similar to that described for definitive endoderm differentiation above, except permeabilization was for 30 mins and primary antibodies Alpha-1-antitrypsin (Bio-Strategy) 1:100 and Human serum albumin (R&D Systems) 1:100 were used. CYP3A4 activity was detected with P450-Glo CYP3A4 Assay with Luciferin-IPA (Promega) following manufacturer guidelines.

### Microfluidic RT-qPCR preparation and analysis

#### RNA extraction

Snap frozen cell pellets had total RNA extracted using ISOLATE II RNA mini kit (Bioline) following manufacturer’s instructions. Briefly, samples were lysed, homogenized and passed through a spin column containing a silica membrane to which the RNA binds. DNase 1 digestion eliminated potential genomic DNA contamination and the preparation was washed to remove impurities such as cellular debris and salts. The purified RNA was eluted with RNase free water and total RNA concentration was determined using Nanodrop ND-1000 Spectrophotometer (Thermo Fisher Scientific). RNA was used either for Microfluidic RT-qPCR or RNA-sequencing.

#### cDNA synthesis and preparation

Total RNA was adjusted to a concentration of 200ng/*μ*l. A 5*μ*l reaction mix was prepared composing of 1*μ*l Reverse Transcription Master Mix (Fluidigm), 3*μ*l of RNase free water and 1*μ*l of RNA and incubated in a thermocycler using the following conditions: 25°C for 5 min, 42°C for 30 min and 85°C for 5 min.

#### cDNA preamplification

3.75*μ*l of preamplification mix (comprising 105.6*μ*l of Preamp MasterMix (Fluidigm), 52*μ*l of 100 *μ*M pooled primer and 237.6*μ*l DNAse/RNAse free water) and 1.25*μ*l of cDNA sample was added into each well of a 96 well plate and incubated as follows: 95°C for 2 min then 10 cycles of 95°C for 15 sec and 60°C for 4 min.

#### cDNA clean-up

cDNA clean-up was performed by adding 2*μ*l of the following mix: 168*μ*l DNase free water, 24*μ*l 10x Exo1 reaction buffer and 48*μ*l Exonuclease I (New England Biolabs), into each well and incubating in a thermocycler for 30 min at 37°C followed by 15 min at 80°C. Samples were diluted 10x with low EDTA TE buffer.

#### Primer and sample set-up

A sample mix was prepared as follows (per 96-well plate): 495*μ*l of 2X SsoFast EvaGreen SuperMix with low ROX (Biorad) and 49.5*μ*l 25X DNA Binding Dye (Fluidigm). 4.95*μ*l sample mix was added with 4.05*μ*l of diluted sample. Primers were prepared in the following mix (per 96-well plate): 450*μ*l 2X Assay Loading Reagent (Fluidigm) and 405*μ*l low EDTA TE buffer. 105*μ*l of primer mix was added with 0.45*μ*l combined forward and reverse primers (Supplementary Table 2). Samples and primer mixes were loaded onto a 96.96 Dynamic Array IFC plate (Fluidigm) and run on the Biomark HD System (Fluidigm).

### CRISPR-KO engineering

#### gRNA design and cloning

Single *S. pyogens* Cas9 gRNA (GCGCCGCGTTTCCAGCGTAC<u>CGG</u>) targeting *MIXL1* exon 1 was designed using Geneious software, (http://www.geneious.com^48^) based on the presence of a canonical NGG PAM (underlined in gRNA sequence) at the target site. Potential off-target sites were identified using Geneious software. gRNA was cloned in Addgene plasmid 62988 following adopted protocol from Ran et al^49^. Oligos used for cloning were:

Forward: 5’ CACCGCGCCGCGTTTCCAGCGTAC

Reverse: 5’ AAACGTACGCTGGAAACGCGGCGC.

#### Nucleofection, clone selection and sequencing

Cells were transfected using a plasmid expressing Cas9 protein and gRNA targeting MIXL1 exon 1 following Amaxa™ 4D Transfection protocol for 20 *μ*l Nucleocuvette® Strip using P3 Primary Cell 4D-Nucleofector® X Kit with program CA-137. After transfection, cells were plated into 10 cm dish coated with hESC-qualified Matrigel (BD Biosciences), prefilled with mTESR1 medium (StemCell Technologies) mixed with 100% CloneR (StemCell Technologies). Twenty-four hours post transfection, cells were puromycin (Thermo Fisher Scientific) selected with concentration of 1*μ*g/ml for the next 48 h. Following puromycin selection media was changed every day and the percentage of CloneR (Stem Cell Technologies) in media was reduced during the next days as single cells were dividing and starting to form individual colonies. Single colonies were picked and transferred individually in a single well of a 96 well plate where they were grown to be split and frozen for further sequencing analysis. Cells were detached using ReLeSR (StemCell Technologies) and clones were frozen as cell aggregates in CryoStor CS10 (StemCell Technologies). Clone selection, screening of the CRISPR/Cas9 clones for editing events and validation of allelic deletions of individual clones was done following protocol from Bauer et al. ^50^for genomic deletions in mammalian cells. The PCR was designed to amplify the sequence flanking the gRNA on exon 1 targeting location with the expected amplicon of 800bp. PCR analysis for the presence of indels were done with primers: Forward: 5’GGAGGGTATAAGTGCGGCC Reverse: 5’CCTCATCTGTGTCTTCTTCCCG.

All PCR reactions were done in 50*μ*l volume using Q5 high fidelity polymerase (NEB) following NEB Q5 high fidelity PCR protocol. In short, PCR reaction mix was made by mixing 100ng of genomic DNA sample from each clone with 10*μ*l of 5x Q5 reaction Buffer, 1*μ*l of 10mM dNTPs, 2.5*μ*l of each (forward and reverse) 10 *μ*M primer, 10*μ*l of 5x Q5 High GC Enhancer, 0.5*μ*l of Q5 Polymerase and topped up to 50*μ*l with H_2_O. The PCR reaction started with initial denaturation of 98°C for 30 sec followed by 34 cycles of 10 sec denaturation at 98°C, annealing at 60°C for 20 sec and extension at 72°C for 20 sec ending with final extension at 72°C for 5 min. The PCR reaction was run on 1.5% agarose gel where expected amplicon of 800bp for each analyzed clone was detected. In total, 43 samples were separately amplified by PCR and analyzed by sequencing for the presence of indels at the exon 1 targeted site. Next, sequenced clones were analyzed for genome editing and indel percentages were calculated via TIDE^51^ using a control chromatogram for comparison. Decomposition windows, left boundaries, and indel ranges were optimized to have the highest alignment possible. After TIDE analysis, 11 clones were selected for validation of biallelic deletion clones for targeted genomic region of exon 1, which was done following standard protocol from Bauer et al.^50^

### C32-iMIXL1 engineering

We overexpressed MIXL1 in the C32 human iPSC line with lentiviral transgenes carrying a doxycycline inducible dSpCas9-VP64, similar to our methods previously published^52^. Briefly, we performed transduction of C32-dCas9-VP64 line with a blasticidin-selectable lentivirus constitutively expressing two of the pre-validated sgRNAs that direct the transcriptional activator upstream of the MIXL1 promoter (Figure S4C). The sgRNA sequences were (+) GAAGAGAGTTCTGTCGCCTG and (+) GCCCAGGCCATGTAAGGCAC. The transduced cell line was clonally derived and selected based on the MIXL1 expression following Doxycycline induction.

### ATAC-seq samples preparation and analysis

#### Cell Preparation

hiPSC were collected at Day 1 of the DE diff protocol (STEMDiff) and were processed following the protocol by Salehin et al^53^ Briefly, 5 x 10^4^ cells were collected at Day 1 of DE differentiation and lysed in cold lysis buffer (10 mM Tris·Cl, pH 7.4, 10 mM NaCl, 3 mM MgCl_2_, 0.1% (v/v) Igepal CA-630). Intact nuclei were separated by centrifugation at 500xg and immediately digested in transposase mix containing 25*μ*l 2x Tagment DNA buffer, 2.5*μ*l Tagment DNA enzyme I (Illumina) and 22.5*μ*l nuclease-free water for 30 min at 37°C. Digested chromatin fragments were then purified using the MinElute PCR Purification Kit (Qiagen), according to manufacturer’s instructions. The fragments of DNA were then pre-amplified by adding 10*μ*L purified DNA sample, 10*μ*L RNase-free water, 2.5*μ*L of each primer (Where each reaction had non-barcoded primer “Ad1_noMix” and one barcoded primer ’Ad2.1’ - ’Ad2.9’ added) (Supplementary Table 3) and 25*μ*L NEBNext High-Fidelity 2x PCR Master Mix (NEB) and was run under the following conditions: 72°C for 5 min, 98°C for 30 sec and then 5 cycles of 98°C for 10 sec, 63°C for 30 sec and 72°C for 1 min. The number of additional cycles to run was calculated by running a RT-qPCR side reaction - a reaction mixture containing 5*μ*L of the pre-amplified PCR product, 3.9*μ*L nuclease-free water, 0.25*μ*L of each primer, 0.6*μ*l 25x SYBR Green and 5*μ*L NEBNext High-Fidelity 2x PCR Master Mix was run under the following conditions: 98°C for 30 sec and then 20 cycles of 98°C for 10 sec, 63°C for 30 sec and 72°C for 1 min. The linear fluorescence versus cycle number was plotted and the cycle number (N) required to reach one-third of the maximum relative fluorescence was determined. The final amplification reaction (the remaining 45*μ*l pre-amplified PCR product) was run under the following conditions: 98°C for 30 sec and then N cycles of 98°C for 10 sec, 63°C for 30 sec and 72°C for 1 min. Amplified samples were then purified using AMPure XP magnetic beads (Beckman Coulter) to remove small fragments and primer-dimers less than 100 bp long (1.3x beads) and large fragments (0.5x beads) using a Dynamag-2 magnet (Thermo Fisher Scientific).

To determine the integrity, fragment size and concentration, the DNA library was analyzed using the Agilent HSD5000 ScreenTape System (Agilent). Libraries were then 101 bp paired-end sequenced on an Illumina HiSeq 4000 (Illumina).

#### Data analysis

ATAC-seq reads were processed using the alignment and filtering functions of the PreNet pipeline^54^. Paired-end reads were mapped to the hg38 genome using bowtie2^55^, allowing for local mapping, a maximum insert size of 2000 bp and a maximum of 4 multimapping hits (–local -X 2000 -k 4). Multimapping reads were allocated using ’assignmultimappers.py’ from the ENCODE ATAC-seq pipeline (https://github.com/ENCODE-DCC/atac-seq-pipeline/tree/master/src). Reads with MAPQ < 30 were excluded and only unique, paired reads that aligned outside blacklisted regions^56^ were used for subsequent analyses. Filtering steps were performed using samtools^57^ and sambamba^58^. Qualifying reads were then converted to pseudo-single end reads and peaks were detected using MACS2^59^, with BED input files and reads shifted by -100 bp and extended to 200bp to capture Tn5 transposase events: -f BED – shift -100 –extend 200 -q 0.05. Biological replicates were analyzed individually and then consensus peak list was created to include only peaks appearing in at least two of the three replicates. Accessibility within a pooled consensus peak list was estimated by quantifying Tn5 events in each of the biological replicates using featureCounts^60^. The DESeq2^61^ package within R was used to identify differentially accessible regions. The regions were filtered for adjusted P-value < 0.05 and an absolute Log2(Fold change) > 1. Coverage tracks were created using the bamCoverage function of deepTools^62^ (–normalization RPKM -bs 10) and visualized within Integrative Genomics Viewer.^63^ findMotifsGenome.pl from HOMER^64^ was used to identify over enriched motifs, between 6 bp and 12 bp in size, within regions of differential accessibility using a repeat masked version of the hg38 sequence (-mask -len 6,7,8,9,10,11,12). Coverage tracks summarizing and combining biological replicates were created using WiggleTools^65^ to quantify the mean coverage per 10 bp bin. These tracks were used for heatmap visualizations created using plotHeatmap from deeptools.

### Micropattern preparation

#### Micropattern chip fabrication

Micropattern chip fabrication was conducted using the protocol of Lee et al.^66^, with specific modification and optimization. In brief, coverslips were sonicated in 70% ethanol for 15 min and in deionized water for 15 min. The clean coverslips were sequentially incubated in 0.5% (3-aminopropyl)triethoxysilane (APTS) (Sigma-Aldrich) for 3 min, 0.5% glutaraldehyde (Sigma-Aldrich) for 30 min. After air drying, the coverslips were deposited on a 20*μ*L drop made of 10% acrylamide (Merck), 0.87% bisacrylamide (Sigma-Aldrich), 0.1% ammonium persulfate (Sigma-Aldrich) and 0.1% N,N,N’,N’-Tetramethylethylenediamine (Sigma-Aldrich), to make the gel at a stiffness of 100KPa. After the stiffness droplet was semi-solidified, the whole system was submerged into 70% ethanol, resulting in a smooth polyacrylamide gel forming. Gelled coverslips were sequentially coated with 64% hydrazine hydrate (Fisher Scientific) for 1 h and 2% glacial acetic acid (Sigma-Aldrich) for 1 h. To generate polydimethylsiloxane (PDMS) stamp, SYLGARD™ 184 Silicone Elastomer Curing Agent and Base (Dow) were mixed at a 1:10 ratio before loading to the stamp mold, provided by the Kilian Lab at the University of New South Wales. Next, the solidified PDMS stamp was coated with 25*μ*g/mL vitronectin (Life Technologies) and 3.5 mg/mL sodium periodate (Sigma-Aldrich) for 1 h. After air-drying the stamp, patterned vitronectin was stamped onto the gelled coverslip at 0.343N for 1 min. Stamped gels were stored overnight in PBS + 1% Penicillin-Streptomycin at 4°C.

#### Germ layer differentiation on micropatterns and analysis

Differentiation protocol was adapted from Warmflash et al.^42^ Since the micropatterned chip generation required many hands-on manipulations, all culture media were supplemented with 1% Penicillin-Streptomycin. hiPSCs were seeded as single cells to micropattern chip at a density of 2.5×10^5^ cells/cm^2^ with 10*μ*M Y-27632 ROCK inhibitor in mTeSR Plus supplemented 1% Penicillin-Streptomycin into a total volume of 1mL per well of a 24-well plate. At about 80% confluency, germ layer differentiation was induced by adding 50 ng/mL BMP4 (R&D Systems) in mTeSR1. The cells grown on micropatterns were washed with PBS, fixed in 4% paraformaldehyde in PBS at RT for 20 min. They were washed with PBS twice and then permeabilised with 0.1% Triton X-100 (Sigma-Aldrich) in dPBS (Gibco) (PBST) at RT for 1 h. The cells were blocked with 3% bovine serum albumin (Sigma-Aldrich) in PBST at room temperature for 1 h. They were incubated with primary antibody at 4°C overnight (MIXL1 (Abcam) 1:50, T/Brachyury (Santa Cruz) 1:50, FOXA2 (Abcam) 1:300, SOX17(R&D Systems) 1:20). Cells were washed with PBST three times, then incubated with corresponding secondary antibodies at RT for 1 h. The cell nuclei were stained with DAPI (1 *μ*g/ml) (Thermo Fisher Scientific) in dPBS for 10 min at RT, and then washed twice more with PBS. Cells were mounted with Fluoromount-G (Thermo Fisher Scientific). Micropatterned coverslips were imaged on Zeiss AiryScan LSM880 confocal microscope.

All image analysis was performed using a custom macro. The nuclei from micropatterned images taken were segmented using the StarDist method^67^ via Fiji software^68^ using default parameters (except probability/score threshold = 0.7) and the versatile (fluorescent nuclei).pb model. R software was used with a custom script where target protein immunofluorescence was normalized to the DAPI intensity of the same cell.

### Bulk RNA sequencing analysis

#### Cell preparation and library prep

lllumina RNA Library prep was performed by GENEWIZ (https://www.genewiz.com/Public/Services/Next-Generation-Sequencing). Samples at Day 1 of DE differentiation (1 - 20*μ*g RNA) were run on HiSeq4000 with a read depth of 20M paired end reads (2x 150PE).

#### Data pre-processing

Details of the procedure can be found in Aryamanesh and colleagues ^69^.

#### Statistical Analysis

Details of the procedure can be found in Aryamanesh and colleagues ^69^.

### Single-cell RNA sequencing

#### Cell preparation

hiPSCs were dissociated using StemPro Accutase (Life Technologies). Single cell suspensions were passed through 40*μ*m cell strainer (Corning) and concentration was adjusted to 1000 cells/*μ*L. Suspensions were loaded in single-cell-G Chip (10X Genomics) for target output of 10,000 cells per sample. Single-cell droplet capture was performed on the Chromium Controller (10X Genomics). cDNA library preparation was performed in accordance with the Single-Cell 3’ v 3.0 or v3.1 protocol. Libraries were evaluated for fragment size and concentration using Agilent HSD5000 ScreenTape System (Agilent).

Samples were sequenced on an Illumina HiSeq4000 instrument according to manufacturer’s instructions (Illumina). Sequencing was carried out using 2×150 paired-end (PE) configuration with a sequencing depth of 20,000 reads per cell. Sequencing was performed by GENEWIZ (Azenta Life Sciences).

#### Data pre-processing

Details of the procedure can be found in Aryamanesh and colleagues ^69^

#### Statistical Analysis

Details of the procedure can be found in Aryamanesh and colleagues ^69^

## Quantification and Statistical Analysis

Statistical analysis was performed using R software. The type of statistical test performed, the meaning of dispersion and precision measurements as well as the significance of each experiment is indicated in the corresponding figure, figure legends and/or in the method details. Outliers have been omitted to facilitate visualization. For micropattern quantification, images were taken from 3 to 10 micropatterns within a coverslip. For microfluidic PCR, three biological replicates were collected for each cell line and timepoint.

## Supplementary Figures

**Figure S1:**
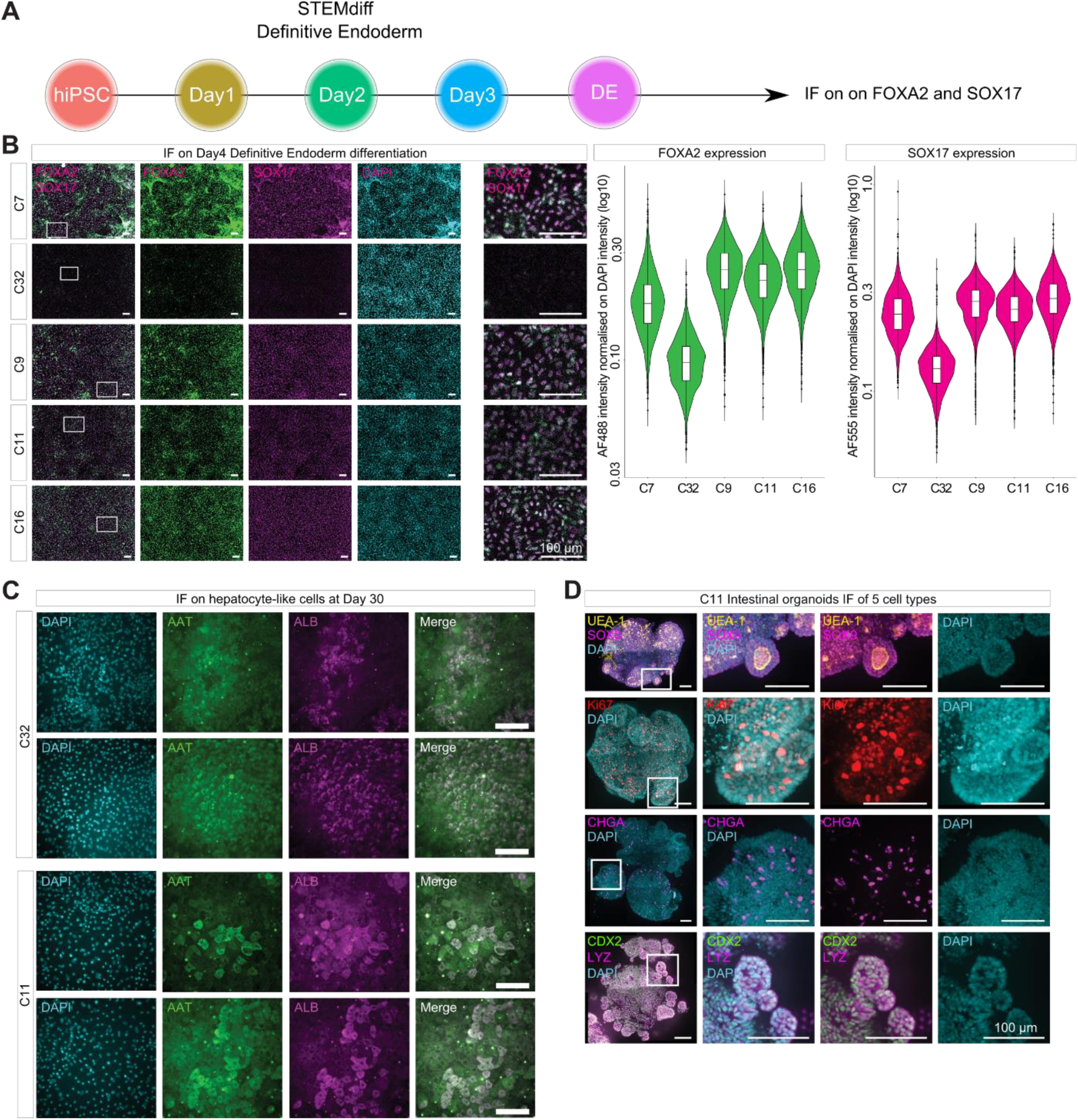
**A)** Definitive endoderm (DE) differentiation protocol used to perform FOXA2 and SOX17 immunofluorescence staining; **B)** Visualization of immunofluoresence of FOXA2 (green) and SOX17 (magenta) in Day 4 cultures (confocal images: column 1-4 from the right, zoom-in images: column 5) and quantification of expression of FOXA2 and SOX17 by fluorescence intensity in violin plots. **C)** Immunofluoresence for detecting AAT (green) and ALB (magenta) in the hepatocytes. B, C: Scale bar: 100µm.

**Figure S2:**
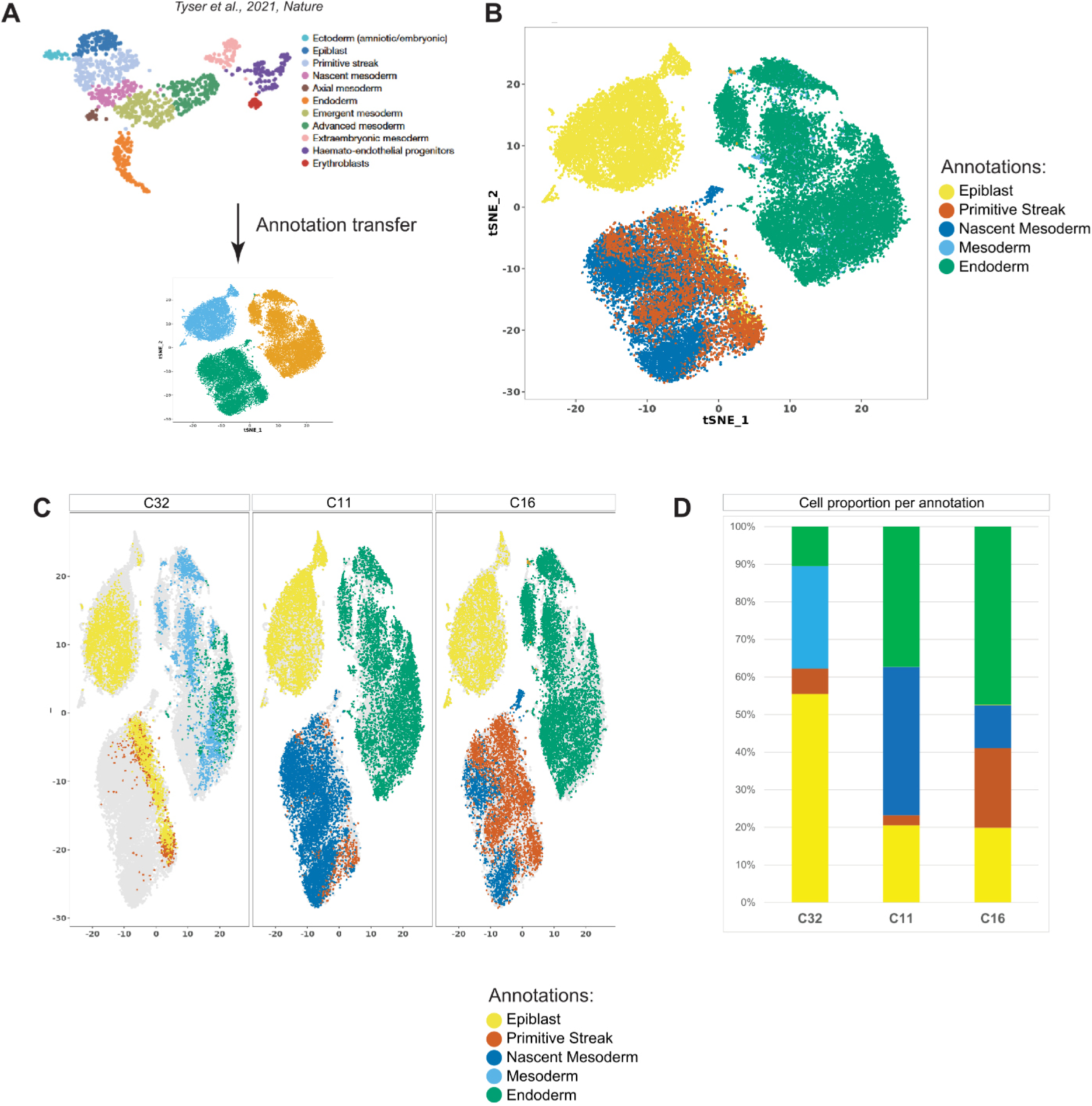
**A)** Schematic of the annotation transfer from Tyser et. al. human post-implantation embryo dataset to our scRNAseq dataset from C32, C11 and C16 at Day 0 (pluripotency), Day 1 (early germ layer differentiation) and Day 4 (definitive endoderm) of differentiation; **B)** Result of the annotation transfer applied on our scRNA-seq dataset; **C)** The same results from S2B separated into the individual C32, C11 and C16 cell lines; **D)** Stacked barplot representing the proportion of each cell type for C32, C11 and C16 cell lines.

**Figure S3:**
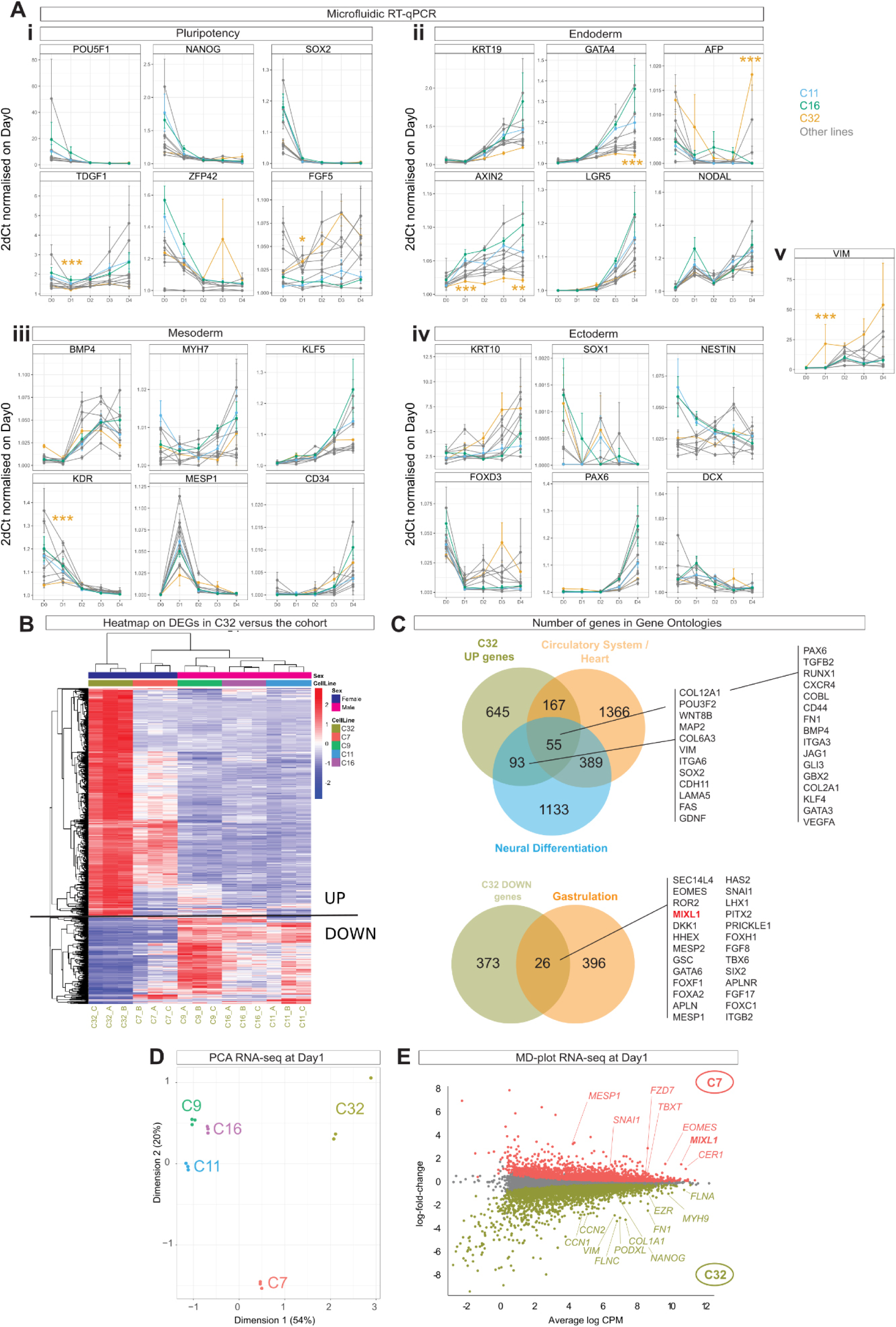
**A)** Microfluidic RT-qPCR data of gene expression by time course during DE differentiation. C16, C11 and C32 are highlighted. p.value: significant differences between C32 versus at least C11 and C16 where * < 0.05, ** < 0.01, *** < 0.001. **i)** Pluripotency-, **ii)** Endoderm-, **iii)** Mesoderm-, **iv)** Ectoderm-related genes and **v)** Vim. **B)** Heatmap of differentially expressed genes (DEGs) of C32 cell line versus ten other cell lines based on bulk RNAseq data. **C)** Venn diagram of upregulated and down-regulated DEGs annotated by relevant ontologies based on bulk RNAseq data. **D)** PCA presenting the bulk RNAseq data of Day 1 samples. **E)** DEGs between C32 and C7 based on Day 1bulk RNAseq data. DEG at p.value < 0.05.

**Figure S4:**
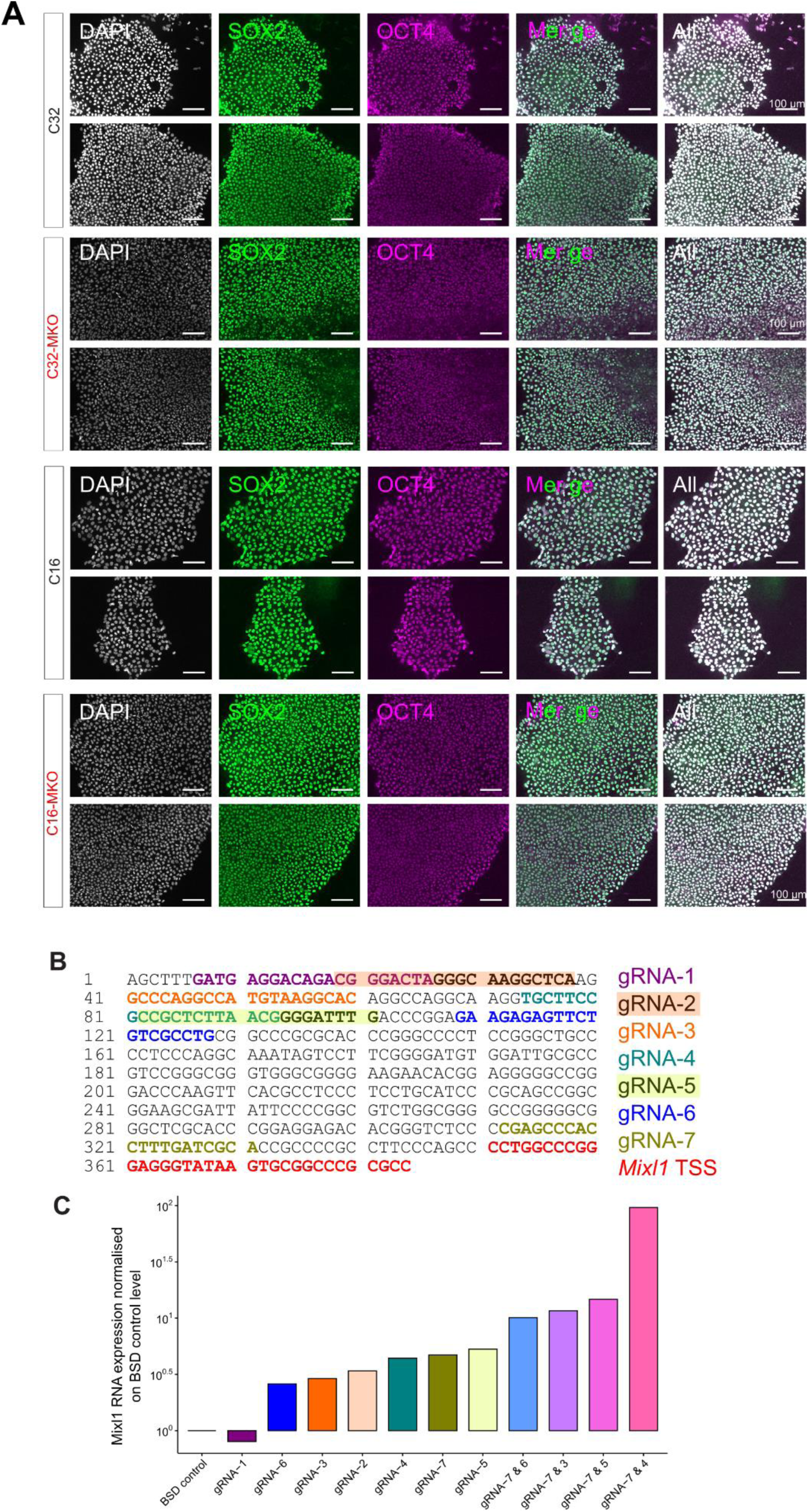
**A)** Expression of SOX2 (green) and OCT4 (magenta) on two colonies each of C32, C32-MKO, C16 and C16-MKO lines. Scale bar: 100µm. **B)** Seven guide RNA predicted by Benchling, targeted to the genomic region of 350bp upstream of *Mix/1* TSS. **C)** RT-qPCR analysis of *Mix/1* expression in HEK cells transfected with single guides or guides in tandems.

**Figure S5:**
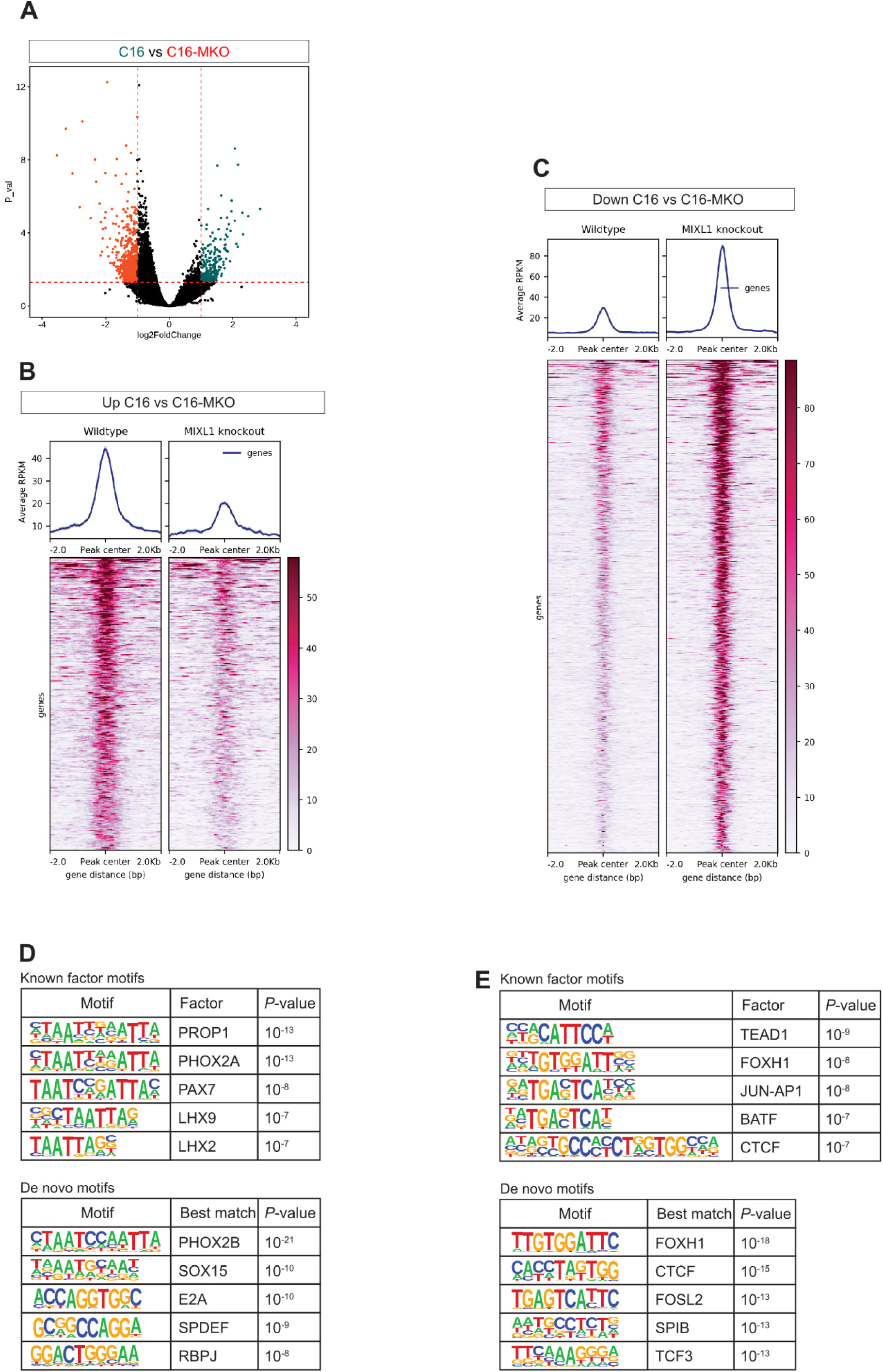
**A)** Volcano plot presenting the Differentially Accessible Chromatins (DACs) region between C16 and C16-MKO lines. **B, C)** Heatmap showing ATAC-seq signal in C16 and C16-KO lines for DACs that are **B)** more accessible in C16 and **C)** in C16-MKO lines. **D, E)** Motifs enriched in DACs more accessible in **D)** C16 and **E)** C16-MKO lines.

**Figure S6:**
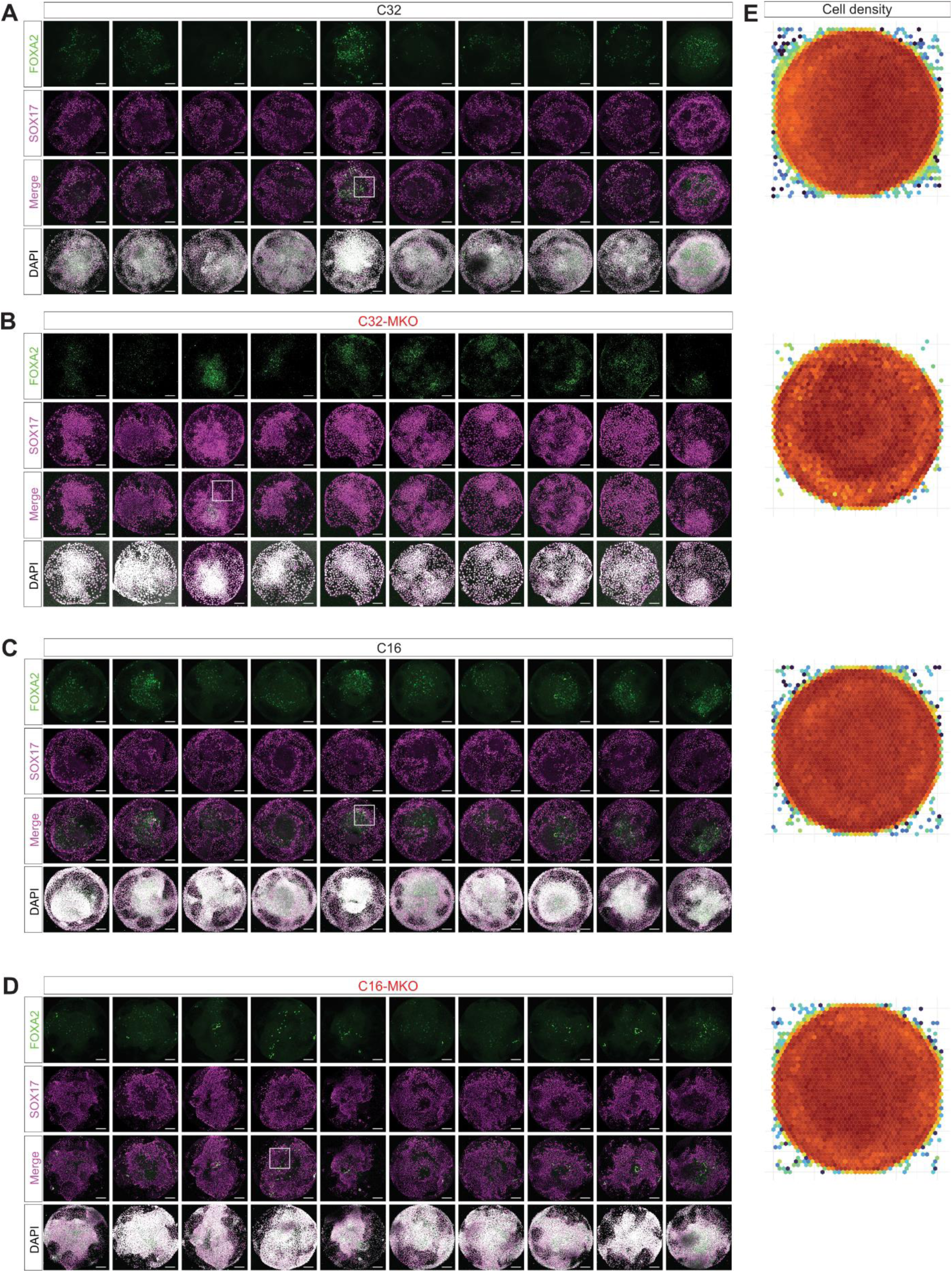
Micropatterns showing expression of FOXA2 and SOX17 at 48h post BMP4 treatment of **A)** C32, **B)** C32-MKO, **C)** C16 and **D)** C16-MKO Lines. N=10 for each line. **E)** The average cell density of all ten micropatterns analyzed for each line.

**Figure S7:**
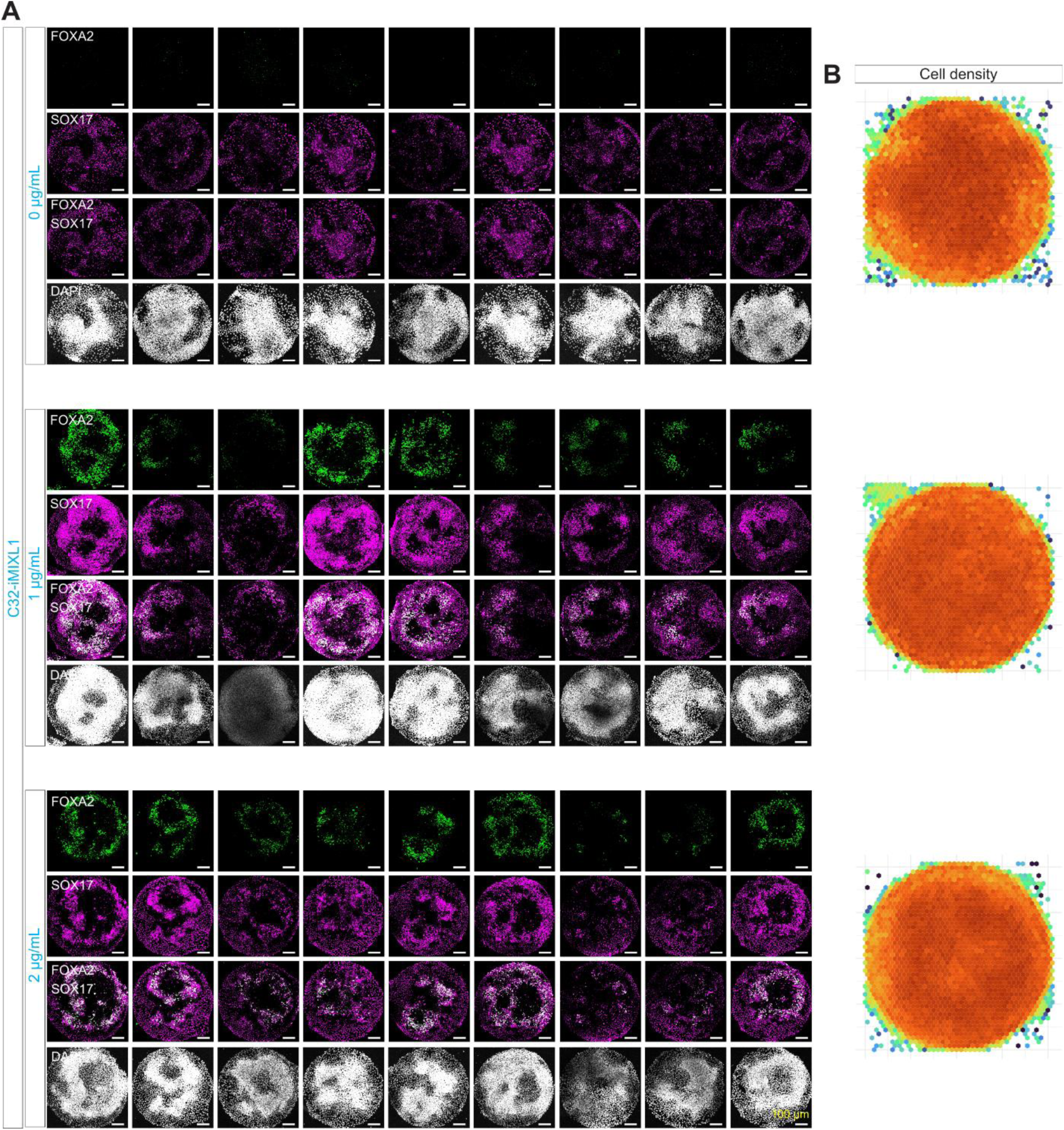
**A)** Micropatterns showing expression of FOXA2 and SOX17 at 48h post BMP4, in C32-iMXL1 line induced with 0, 1 and 2µg/mL of Doxycycline. N=9 for each dose. **B)** The average cell density for micropatterns of C32-iMXL1 line induced with 0, 1 and 2µg/mL of doxycycline.

## Key resources table

### Key resources table

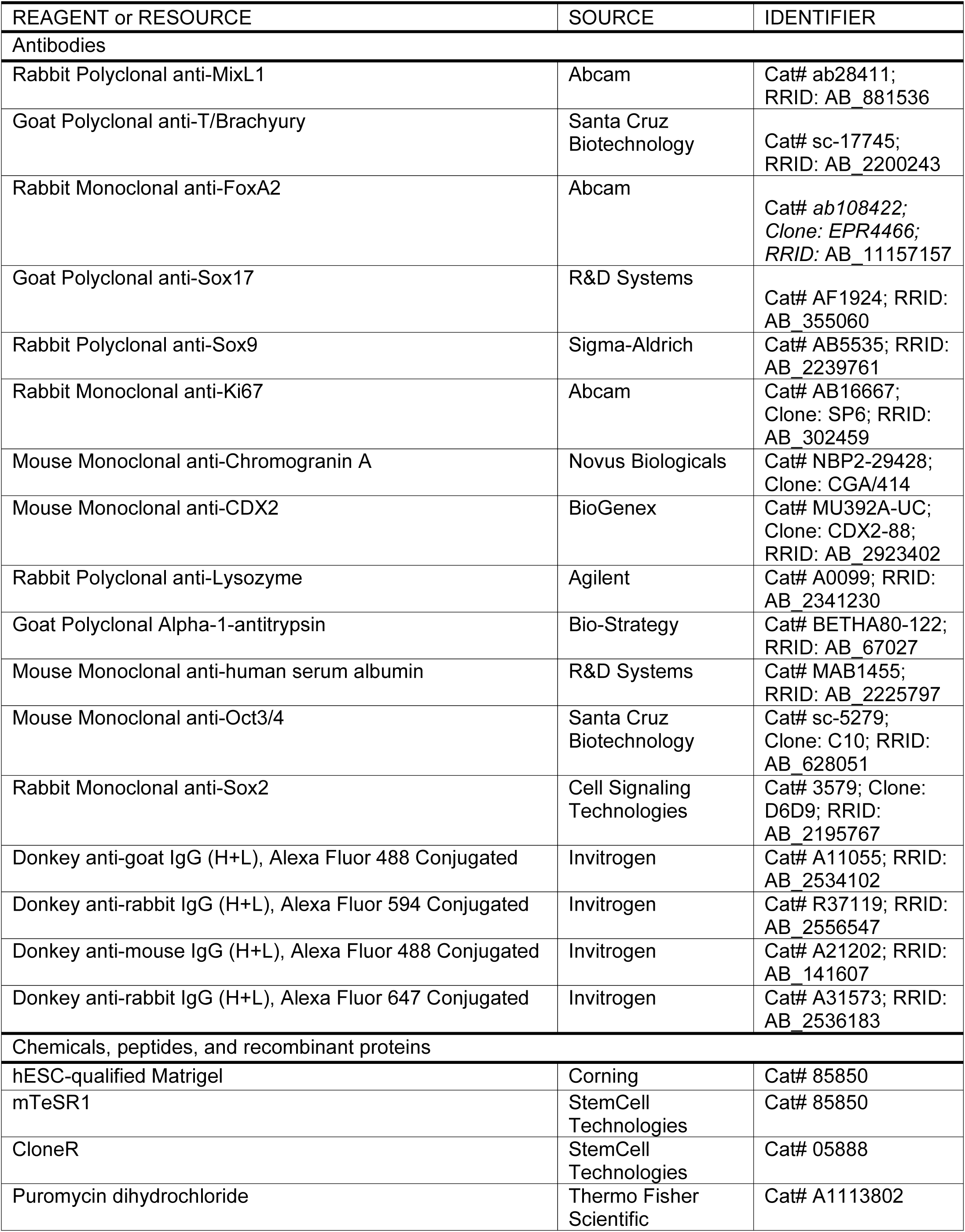
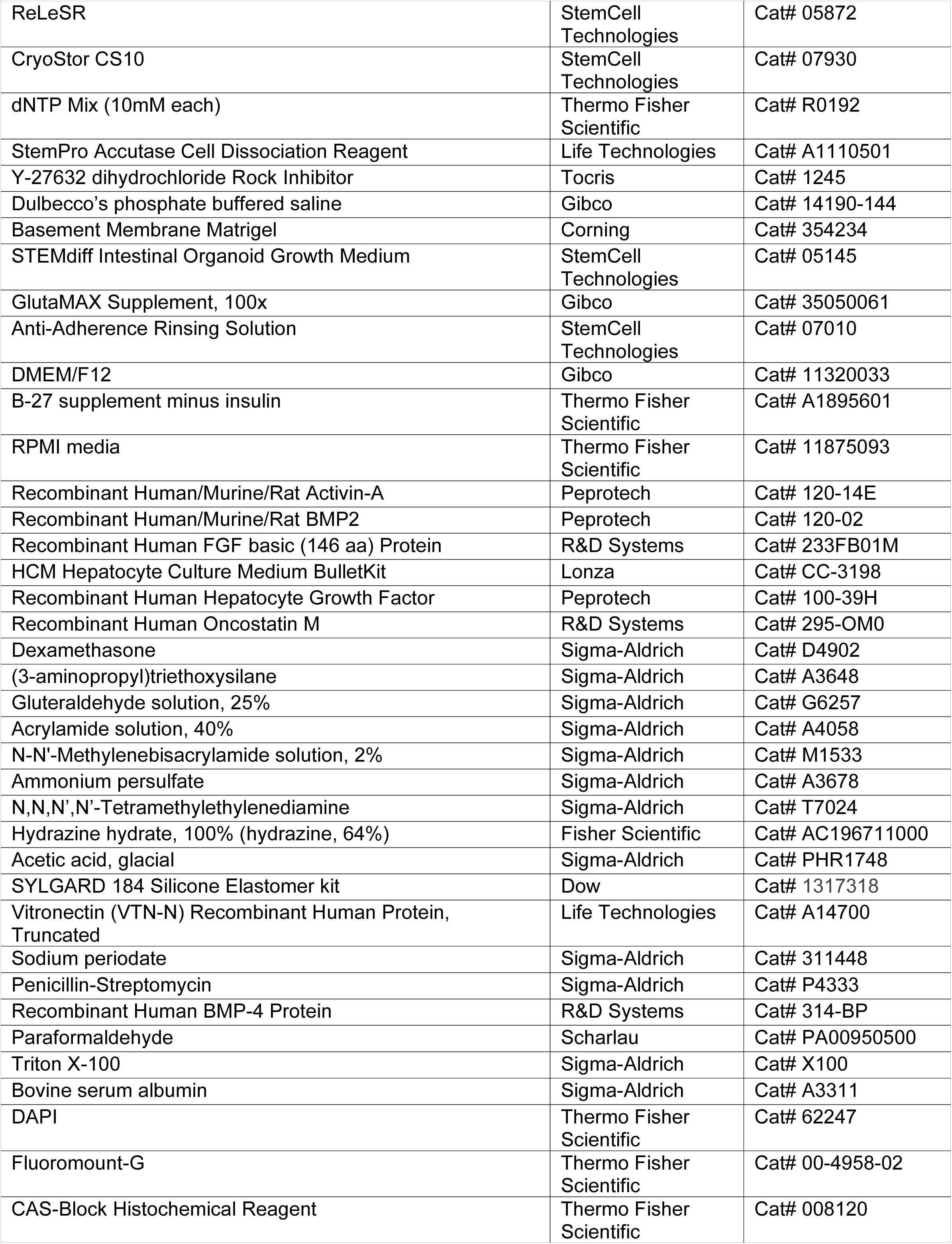
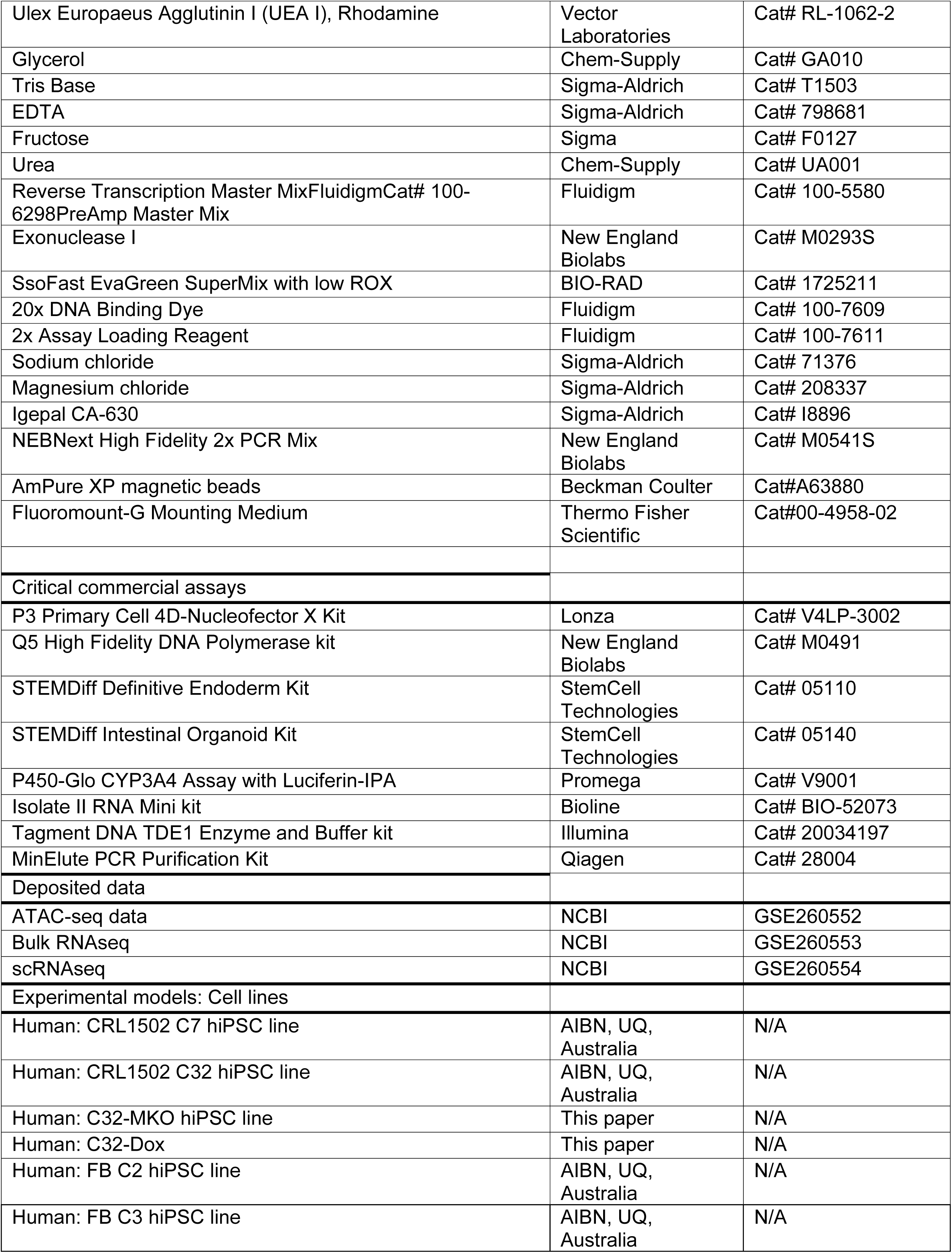
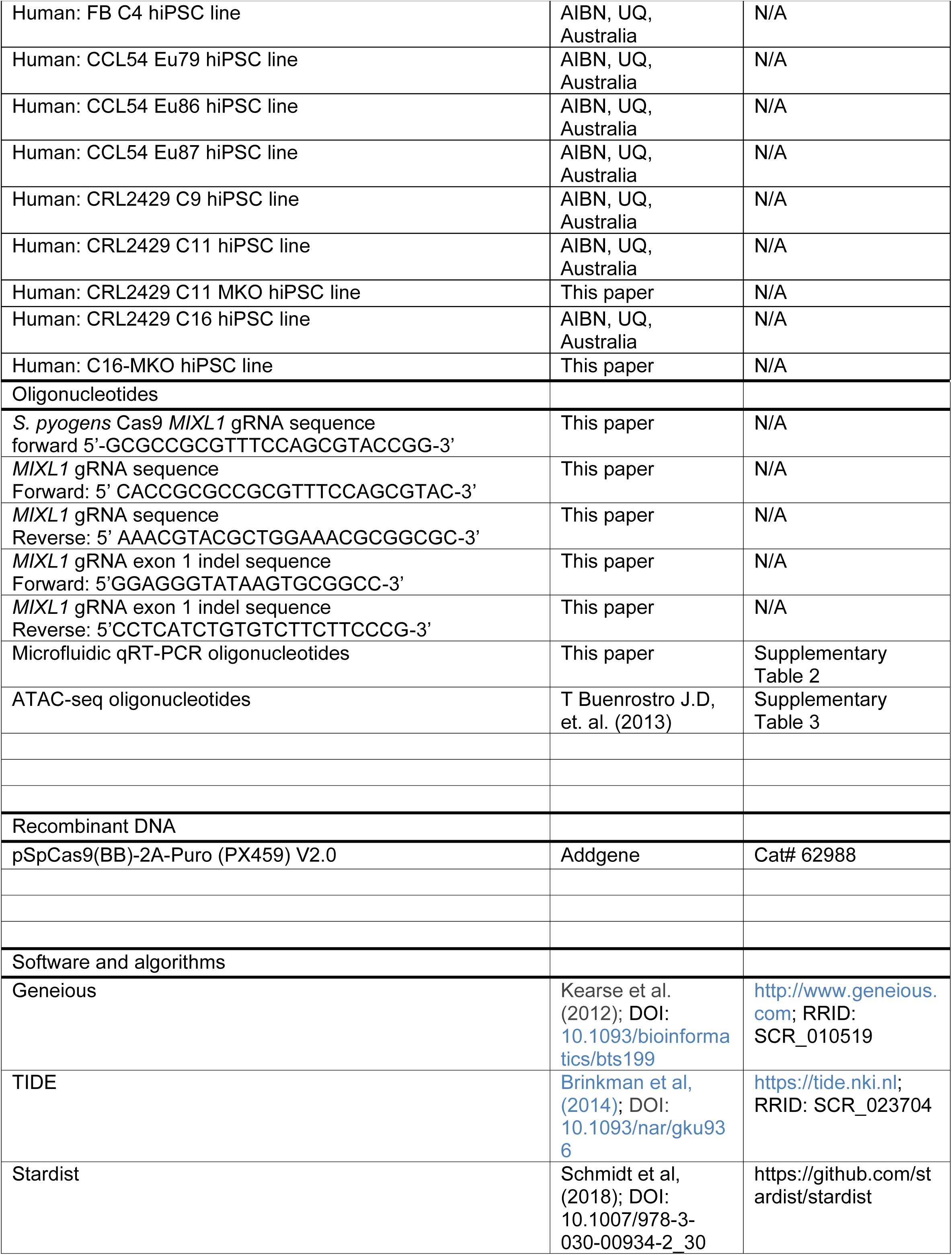
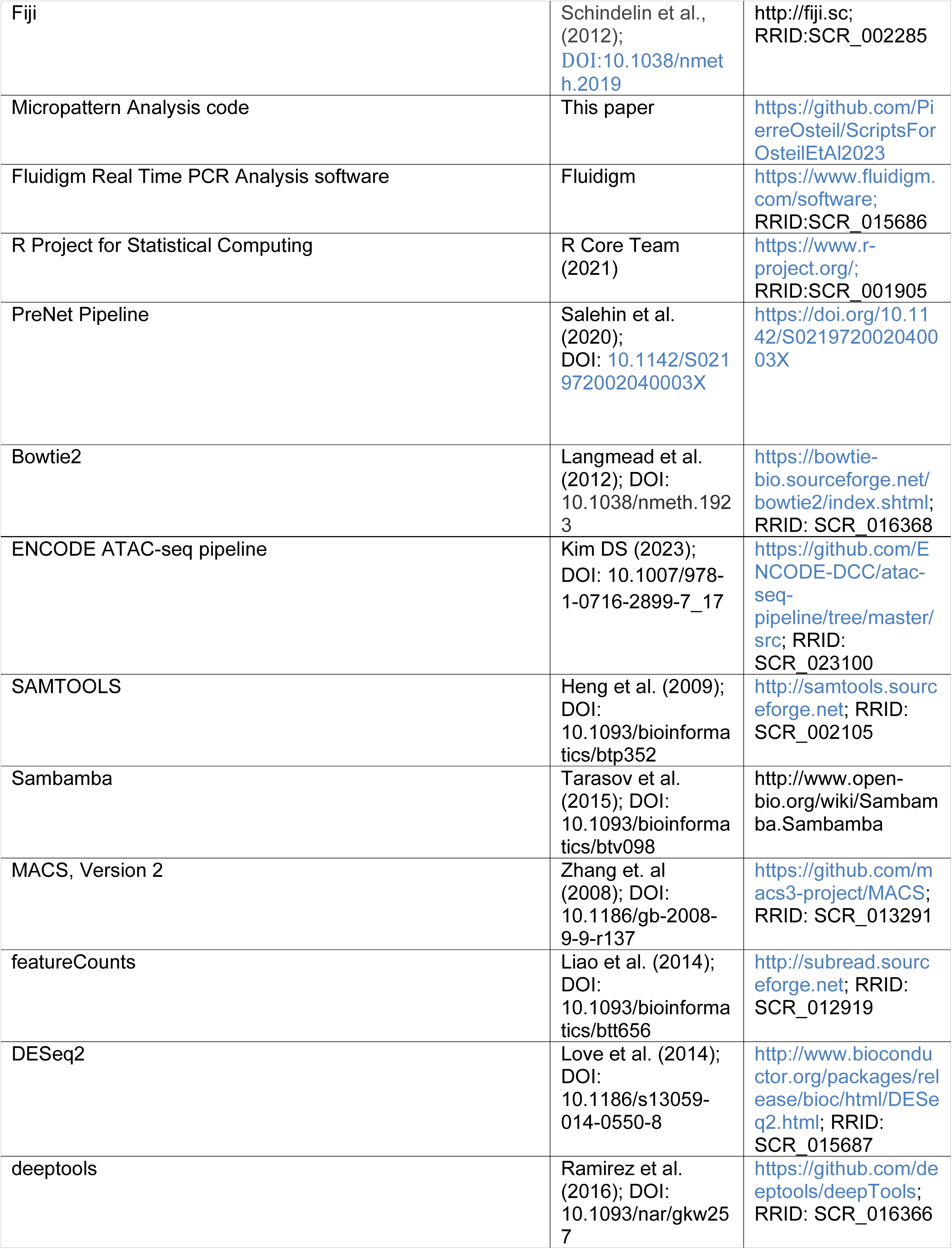
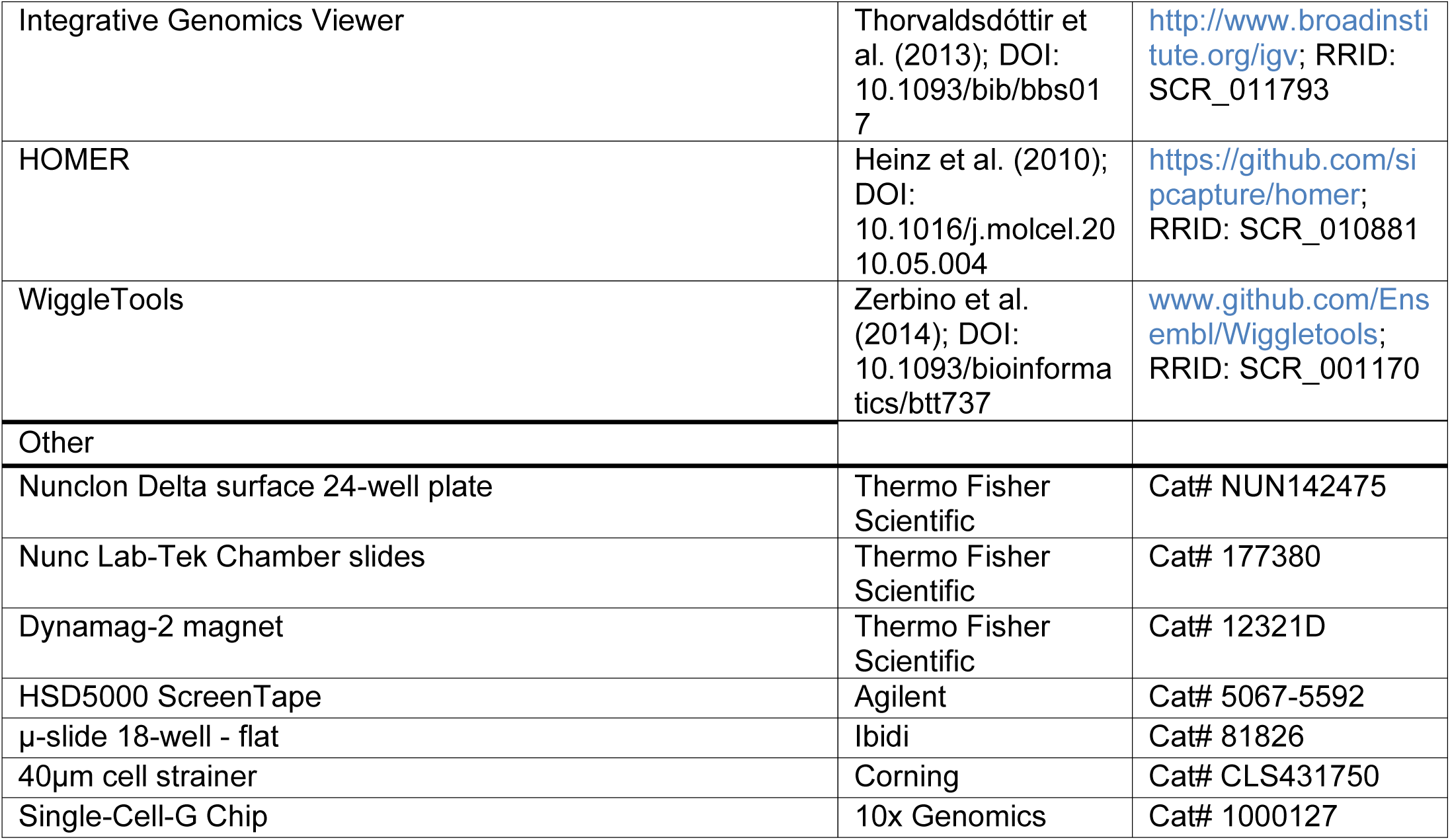

