## Supplementary material for "MIXL1 Activation in Endoderm Differentiation of Human Induced Pluripotent Stem Cells": Cell line meatadata

| **Name of iPSc line** | **Karyotype** | **Pluripotency markers** | **Transgene detection** | **Teratoma** | **Gender** | **Age** | **Reprogramming method** | **Cell type of origin** |
| --- | --- | --- | --- | --- | --- | --- | --- | --- |
| FB C3 | y | y | y | y | F | 25+ | Episomal | Fibrobalst |
| FB C2 | Y | Y | Y | ? | F | 25+ | Episomal | Fibrobalst |
| FB C4 | Y | Y | Y? | Y? | F | 25+ | Episomal | Fibrobalst |
| CRL2429 c11 | Y | Y | Y | Y | M | newborn | Episomal | Fibrobalst  (foreskin) |
| CRL2429 C16 | Y | Y | Y | Y | M | newborn | Episomal | Fibrobalst  (foreskin) |
| CRL2429 C9 | Y | Y | Y | Y | M | newborn | Episomal | Fibrobalst  (foreskin) |
| CRL1502 C32 | Y | Y | Y | Y | F | 12 wpc | Episomal | Fibrobalst |
| CRL1502 C29 | Y | Y | Y | Y | F | 12 wpc | Episomal | Fibrobalst |
| CRL1502 C7 | Y | Y | Y | Y | F | 12 wpc | Episomal | Fibrobalst |
| CCL54 Eu79 | Y | Y | Y | Y | M | 25+ | Episomal | Fibrobalst |
| CCL54 Eu86 | Y | Y | Y | ? | M | 25+ | Episomal | Fibrobalst |
| CCL54 Eu87 | Y | Y | Y | Y | M | 25+ | Episomal | Fibrobalst |
