## Supplementary material for "MIXL1 Activation in Endoderm Differentiation of Human Induced Pluripotent Stem Cells": Microfluidic oligos

| Gene | Forward | Reverse | Ref GetPrime |
| --- | --- | --- | --- |
| Zic3 | AAACTGGTCAACCACATCC | TGAAAGGTTTCTCACCTGTG | [2004931](http://bbcftools.epfl.ch/getprime/primers/details?ensembl_version=81&primer_id=2004931) |
| Dppa4 | CAGTAAAGTGCTCTGCCCT | GAAGGTAATGGAGGGATTGGT | [2318106](http://bbcftools.epfl.ch/getprime/primers/details?ensembl_version=81&primer_id=2318106) |
| Fgf5 | GGGATTGTAGGAATACGAGGA | GAAGCGTCCCTTTAAAGGG | [2362133](http://bbcftools.epfl.ch/getprime/primers/details?ensembl_version=81&primer_id=2362133) |
| Klf2 | CATTCCAGTGCCATCTGTG | ACATGTGCCGTTTCATGTG | [1876091](http://bbcftools.epfl.ch/getprime/primers/details?ensembl_version=81&primer_id=1876091) |
| Klf4 | CCCACACTTGTGATTACGC | GGTAAGGTTTCTCACCTGTG | [1936758](http://bbcftools.epfl.ch/getprime/primers/details?ensembl_version=81&primer_id=1936758) |
| Nanog | CAGAGACAGAAATACCTCAGC | GGTCTTCACCTGTTTGTAGC | [2249106](http://bbcftools.epfl.ch/getprime/primers/details?ensembl_version=81&primer_id=2249106) |
| Pou5f1 | GAGAAGGATGTGGTCCGAG | TCCTCTCGTTGTGCATAGTC | [2243084](http://bbcftools.epfl.ch/getprime/primers/details?ensembl_version=81&primer_id=2243084) |
| Prdm14 | CAGCTCTGGATCTGATTCTC | GTCTGCATGAGGCATAGAC | [1865525](http://bbcftools.epfl.ch/getprime/primers/details?ensembl_version=81&primer_id=1865525) |
| Sox2 | CGGATTATAAATACCGGCCC | GTGTACTTATCCTTCTTCATGAGC | [2067242](http://bbcftools.epfl.ch/getprime/primers/details?ensembl_version=81&primer_id=2067242) |
| Zfp42 | TCAAATAACCTGAAAGCCCA | CCTCTTGTTCATTCTTGTTCGT | [1866758](http://bbcftools.epfl.ch/getprime/primers/details?ensembl_version=81&primer_id=1866758) |
| T | CTGCTTATCAGAACGAGGAG | GTGATCACTTCTTTCCTTTGC | [1947250](http://bbcftools.epfl.ch/getprime/primers/details?ensembl_version=81&primer_id=1947250) |
| Mixl1 | CGTACCCCGACATCCACTT | GCCTGTTCTGGAACCATACCT | [1787479](http://bbcftools.epfl.ch/getprime/primers/details?ensembl_version=81&primer_id=1787479) |
| Wnt3a | AGATTGGCATCCAGGAGTG | CTCCCTGGTAGCTTTGTCC | [1951111](http://bbcftools.epfl.ch/getprime/primers/details?ensembl_version=81&primer_id=1951111) |
| Nodal | GCATACATCCAGAGTCTGCT | CACATACAGCATGCTCAGC | [2117271](http://bbcftools.epfl.ch/getprime/primers/details?ensembl_version=81&primer_id=2117271) |
| Lhx1 | AATGCAACCTGACCGAGAAG | TACCGAAACACCGGAAGAAG | [1963413](http://bbcftools.epfl.ch/getprime/primers/details?ensembl_version=81&primer_id=1963413) |
| Eomes | ACAGGAGATTTCATTCGGG | TTGTAAGACTATCATCTGGGTG | [2025268](http://bbcftools.epfl.ch/getprime/primers/details?ensembl_version=81&primer_id=2025268) |
| Gsc | GAGGAGAAAGTGGAGGTCTG | CTCCGACTCCTCTGATGAG | [1785151](http://bbcftools.epfl.ch/getprime/primers/details?ensembl_version=81&primer_id=1785151) |
| Tdgf1 | AGAGATGACAGCATTTGGC | TTTAGCTCCTTACTGTGCTG | [2112793](http://bbcftools.epfl.ch/getprime/primers/details?ensembl_version=81&primer_id=2112793) |
| Axin2 | CCTGGCTCCAGAAGATCAC | GTAAGTGACAACCAACTCACTG | [2052559](http://bbcftools.epfl.ch/getprime/primers/details?ensembl_version=81&primer_id=2052559) |
| Tbx6 | TACCAGAACCCACAGATCAC | TTACAGTTTCTGCCGTTCTC | [2114104](http://bbcftools.epfl.ch/getprime/primers/details?ensembl_version=81&primer_id=2114104) |
| Foxa2 | GAGTTAAAGTATGCTGGGAGC | GTTCATGTTGCTCACGGAG | [2003365](http://bbcftools.epfl.ch/getprime/primers/details?ensembl_version=81&primer_id=2003365) |
| Sox17 | TTCATGGTGTGGGCTAAGG | TGTGCAGGTCTGGATTCTG | [1824267](http://bbcftools.epfl.ch/getprime/primers/details?ensembl_version=81&primer_id=1824267) |
| Sox7 | GAACGCCTTCATGGTTTGG | CTTTCCCAGCATCTTGCTG | [2098878](http://bbcftools.epfl.ch/getprime/primers/details?ensembl_version=81&primer_id=2098878) |
| Krt19 | GCGAAGCCAATATGAGGTC | TTCAATTCTTCAGTCCGGC | [2067764](http://bbcftools.epfl.ch/getprime/primers/details?ensembl_version=81&primer_id=2067764) |
| AFP | GCTTACACAAAGAAAGCCC | TAATAATGTCAGCCGCTCC | [1842428](http://bbcftools.epfl.ch/getprime/primers/details?ensembl_version=81&primer_id=1842428) |
| Cdx2 | TACATCACCATCCGGAGGA | TTCCTCTCCTTTGCTCTGC | [1832797](http://bbcftools.epfl.ch/getprime/primers/details?ensembl_version=81&primer_id=1832797) |
| Shh | AAAAGCTGACCCCTTTAGCC | GATGTCGGGGTTGTAATTGG | [1867823](http://bbcftools.epfl.ch/getprime/primers/details?ensembl_version=81&primer_id=1867823) |
| Hhex | TCCCAGGAAGACCTTGAATC | CAATGCCAGTGGTCATCATC | [2317590](http://bbcftools.epfl.ch/getprime/primers/details?ensembl_version=81&primer_id=2317590) |
| Pdx1 | AAGTCTACCAAAGCTCACG | CCTTCTCCAGCTCTAGCAG | [1806690](http://bbcftools.epfl.ch/getprime/primers/details?ensembl_version=81&primer_id=1806690) |
| Acvr1b | CCTTCTACTGCCTGAGCTC | CTCCTTGAGGTGACCACTG | [2001430](http://bbcftools.epfl.ch/getprime/primers/details?ensembl_version=81&primer_id=2001430) |
| Gata4 | AGATGCGTCCCATCAAGAC | CTGACTGAGAACGTCTGGG | [2013061](http://bbcftools.epfl.ch/getprime/primers/details?ensembl_version=81&primer_id=2013061) |
| Pecam1 | AGAGTACCAGGTGTTGGTG | CTCTTTCTTGTCCAGTGTCAC | [1939011](http://bbcftools.epfl.ch/getprime/primers/details?ensembl_version=81&primer_id=1939011) |
| Hnf4a | TACTCCTGCAGATTTAGCCG | GCATTTCTTGAGCCTGCAG | [1870610](http://bbcftools.epfl.ch/getprime/primers/details?ensembl_version=81&primer_id=1870610) |
| Lgr5 | CTCCGATCGCTGAATTTGG | CTTTATTAGGGATGGCAAAGTGG | [2088320](http://bbcftools.epfl.ch/getprime/primers/details?ensembl_version=81&primer_id=2088320) |
| Dcn | TTGGCTAAGTTGGGATTGAG | TGAAGCTCCCTCAGATGAG | [2101122](http://bbcftools.epfl.ch/getprime/primers/details?ensembl_version=81&primer_id=2101122) |
| Alb | GAAACAAACTGCACTTGTTGAG | GCGAAATCATCCATAACAGCT | [2042866](http://bbcftools.epfl.ch/getprime/primers/details?ensembl_version=81&primer_id=2042866) |
| Hgf | CTATGCAGAGGGACAAAGGA | ACATTGGTCTGCAGTATTCAC | [2030593](http://bbcftools.epfl.ch/getprime/primers/details?ensembl_version=81&primer_id=2030593) |
| Fgf8 | TCTGCATGAACAAGAAGGG | CACAATCTCCGTGAAGACG | [1946527](http://bbcftools.epfl.ch/getprime/primers/details?ensembl_version=81&primer_id=1946527) |
| Nod2 | ATCAGGTTGCCGATCTTCAC | TAGAAGGAAGGCAGCCAATC | [2066660](http://bbcftools.epfl.ch/getprime/primers/details?ensembl_version=81&primer_id=2066660) |
| Cxcr4 | GCCTTACTACATTGGGATCAG | CCCTTGCTTGATGATTTCCA | [1960152](http://bbcftools.epfl.ch/getprime/primers/details?ensembl_version=81&primer_id=1960152) |
| Bmp4 | CCACCACGAAGAACATCTG | ATGCTGCTGAGGTTAAAGAG | [1981442](http://bbcftools.epfl.ch/getprime/primers/details?ensembl_version=81&primer_id=1981442) |
| Col10a1 | GATACCAAATGCCCACAGG | CCTCTTACTGCTATACCTTTACTC | [1916930](http://bbcftools.epfl.ch/getprime/primers/details?ensembl_version=81&primer_id=1916930) |
| Ctsk | CCATCCATAACCTTGAGGCT | TCTTCACTGGTCATGTCCC | [1806453](http://bbcftools.epfl.ch/getprime/primers/details?ensembl_version=81&primer_id=1806453) |
| Gata6 | GACTTGCTCTGGTAATAGCA | CTGTAGGTTGTGTTGTGGG | [1816674](http://bbcftools.epfl.ch/getprime/primers/details?ensembl_version=81&primer_id=1816674) |
| Kdr | GTCACTTTGTGCAAGATACCC | GTAAAGCCCTTCTTGCTGTC | [1936266](http://bbcftools.epfl.ch/getprime/primers/details?ensembl_version=81&primer_id=1936266) |
| Gata2 | CAAGCTGCACAATGTTAACAG | GACTTGTTGGACATCTTCCG | [1820193](http://bbcftools.epfl.ch/getprime/primers/details?ensembl_version=81&primer_id=1820193) |
| Hand1 | CTGAACTCAAGAAGGCGGA | AATCCTCTTCTCGACTGGG | [1871030](http://bbcftools.epfl.ch/getprime/primers/details?ensembl_version=81&primer_id=1871030) |
| Hand2 | CAAGAAGACCGACGTGAAAG | TTCTTGTCGTTGCTGCTCAC | [2037568](http://bbcftools.epfl.ch/getprime/primers/details?ensembl_version=81&primer_id=2037568) |
| Igf2 | GAGACGTACTGTGCTACCC | ATTGGAAGAACTTGCCCAC | [1904145](http://bbcftools.epfl.ch/getprime/primers/details?ensembl_version=81&primer_id=1904145) |
| Smtn | TGGACAAGATGCTGGATCAG | TCTTCCTTTGTCGGAGCTC | [1847185](http://bbcftools.epfl.ch/getprime/primers/details?ensembl_version=81&primer_id=1847185) |
| Klf5 | CTACTGCGATTACCCTGGT | GTGTGAGTCCTCAGGTGAG | [2029387](http://bbcftools.epfl.ch/getprime/primers/details?ensembl_version=81&primer_id=2029387) |
| Mesp1 | GTGACAAGGGACAACTGAC | AGGAACCACTTCGAAGGTG | [1807878](http://bbcftools.epfl.ch/getprime/primers/details?ensembl_version=81&primer_id=1807878) |
| Nkx2-5 | TAAACCTGGAACAGCAGCA | TAGGCACGTGGATAGAAGG | [1858829](http://bbcftools.epfl.ch/getprime/primers/details?ensembl_version=81&primer_id=1858829) |
| Pdgfra | GTCTTCTCACAGGGCTGAG | TGAATTCAGCTGCACAACC | [1866737](http://bbcftools.epfl.ch/getprime/primers/details?ensembl_version=81&primer_id=1866737) |
| Runx1 | CATCGCTTTCAAGGTGGTG | GTAGCATTTCTCAGCTCAGC | [1805614](http://bbcftools.epfl.ch/getprime/primers/details?ensembl_version=81&primer_id=1805614) |
| Myl3 | AAGAAGGATGATGCCAAGG | CTCAATCTTGATCTTGGAAGC | [1882070](http://bbcftools.epfl.ch/getprime/primers/details?ensembl_version=81&primer_id=1882070) |
| Myh7 | TGCAGCAGTTCTTCAACCAC | TCTCGATGAGGTCAATGCAG | [1897494](http://bbcftools.epfl.ch/getprime/primers/details?ensembl_version=81&primer_id=1897494) |
| Cd34 | CCAACAGAACAGAAATTTCCAG | CTCAGTGAAATCTAGGATCCC | [2116431](http://bbcftools.epfl.ch/getprime/primers/details?ensembl_version=81&primer_id=2116431) |
| Cd79a | GAACCGAATCATCACAGCC | CCATCGTTTCCTGAACAGC | [1849875](http://bbcftools.epfl.ch/getprime/primers/details?ensembl_version=81&primer_id=1849875) |
| Ryr2 | CAGGACAGGAATCTTATGTCTG | CTGTTTCCGGAAGCAATCC | [1924479](http://bbcftools.epfl.ch/getprime/primers/details?ensembl_version=81&primer_id=1924479) |
| Comp | CAAGACCAGGATGGAGACG | CATCGTGGTCTGAGTCCTC | [1881734](http://bbcftools.epfl.ch/getprime/primers/details?ensembl_version=81&primer_id=1881734) |
| Ccr5 | ATCTTCTTCATCATCCTCCTGAC | CAAACACAGCATGGACGAC | [1907495](http://bbcftools.epfl.ch/getprime/primers/details?ensembl_version=81&primer_id=1907495) |
| Krt10 | CCCGACGGTAGAGTTCTTTC | AATGGTCTGTGTGAAGGGAGAC | [1877417](http://bbcftools.epfl.ch/getprime/primers/details?ensembl_version=81&primer_id=1877417) |
| Zic1 | CGAACAAAGTTCGGAAGCTC | GGATCCGAGGTAGAGGAAAAG | [2025115](http://bbcftools.epfl.ch/getprime/primers/details?ensembl_version=81&primer_id=2025115) |
| Fabp7 | ACGGAGATTAGTTTCCAGCT | CAGGCTAACAACAGACTTACAG | [1942020](http://bbcftools.epfl.ch/getprime/primers/details?ensembl_version=81&primer_id=1942020) |
| Dcx | GAAGCCATCAAACTGGAGAC | GAAATCATGGAGACAAGTTACCTG | [1830742](http://bbcftools.epfl.ch/getprime/primers/details?ensembl_version=81&primer_id=1830742) |
| Nes | TTCCCTCAGCTTTCAGGAC | GAGCAAAGATCCAAGACGC | [1883450](http://bbcftools.epfl.ch/getprime/primers/details?ensembl_version=81&primer_id=1883450) |
| Olig2 | ACTACATCCTCATGCTCACC | TAGATCTCGCTCACCAGTC | [1910459](http://bbcftools.epfl.ch/getprime/primers/details?ensembl_version=81&primer_id=1910459) |
| Chat | AGGAGCAGTTCAGGAAGAG | TACTCAGACACCCAGTTGG | [1889660](http://bbcftools.epfl.ch/getprime/primers/details?ensembl_version=81&primer_id=1889660) |
| Foxd3 | CTACTACAGGGAGAAGTTCCC | GTTGAGTGAGAGGTTGTGG | [2071972](http://bbcftools.epfl.ch/getprime/primers/details?ensembl_version=81&primer_id=2071972) |
| Pax6 | GAAGCAAGAATACAGGTATGGT | CTGTCTTCTCTGATTCCTCAG | [2026912](http://bbcftools.epfl.ch/getprime/primers/details?ensembl_version=81&primer_id=2026912) |
| Vim | TCCACGAAGAGGAAATCCA | CAGGCTTGGAAACATCCAC | [1791140](http://bbcftools.epfl.ch/getprime/primers/details?ensembl_version=81&primer_id=1791140) |
| Pou4f1 | TTCCAATGAGAGGCCTATGG | TGGATTCCACCTAAGCAAGC | [1984741](http://bbcftools.epfl.ch/getprime/primers/details?ensembl_version=81&primer_id=1984741) |
| Eno1 | CTTTACGTTCACCTCGGTG | CATGGTGAACTTCTAGCCAC | [1900509](http://bbcftools.epfl.ch/getprime/primers/details?ensembl_version=81&primer_id=1900509) |
| Gbx2 | TAACTTCGACAAGGCGGAG | CCTTTGACTCGTCTTTCCC | [1957564](http://bbcftools.epfl.ch/getprime/primers/details?ensembl_version=81&primer_id=1957564) |
| Gfap | ACTCAATGCTGGCTTCAAGG | GCCTTGTTTTGCTGTTCCAG | [2054596](http://bbcftools.epfl.ch/getprime/primers/details?ensembl_version=81&primer_id=2054596) |
| Tubb3 | AGTGTGAAAACTGCGACTGC | GCTGAAGGTGTTCATGATGC | [1944357](http://bbcftools.epfl.ch/getprime/primers/details?ensembl_version=81&primer_id=1944357) |
| Gad1 | GACTCTGGACAGTAGAGGC | AGGTATCGTACGTTGTGGG | [2083258](http://bbcftools.epfl.ch/getprime/primers/details?ensembl_version=81&primer_id=2083258) |
| L1cam | AGTGACAAGTACTTCATAGAGG | CACATCCAGTTCGGTACTG | [2018356](http://bbcftools.epfl.ch/getprime/primers/details?ensembl_version=81&primer_id=2018356) |
| Sox1 | GCCATGGATGAAGGACAAAG | GGTTCAGCGATTGTGTTTCC | [1884560](http://bbcftools.epfl.ch/getprime/primers/details?ensembl_version=81&primer_id=1884560) |
| Otx2 | CCAGACATCTTCATGCGAG | TCGATTCTTAAACCATACCTGC | [1968144](http://bbcftools.epfl.ch/getprime/primers/details?ensembl_version=81&primer_id=1968144) |
| Krt14 | AGAGAAGAACCGCAAGGATG | AATCTCCAGGTTCTGCATGG | [1877723](http://bbcftools.epfl.ch/getprime/primers/details?ensembl_version=81&primer_id=1877723) |
| Grin1 | CCTGTCCATCCTCAAGTCC | GAGTCACATTCCTGATACCGA | [1992840](http://bbcftools.epfl.ch/getprime/primers/details?ensembl_version=81&primer_id=1992840) |
| TH | CCTGGTTCCCAAGAAAAGTG | AAGGCGATCTCAGCAATCAG | [2047462](http://bbcftools.epfl.ch/getprime/primers/details?ensembl_version=81&primer_id=2047462) |
| Gapdh | TCAAGATCATCAGCAATGCC | CGATACCAAAGTTGTCATGGA | [1936698](http://bbcftools.epfl.ch/getprime/primers/details?ensembl_version=81&primer_id=1936698) |
| Actb | AGAAGGATTCCTATGTGGGC | TACTTCAGGGTGAGGATGC | [2013505](http://bbcftools.epfl.ch/getprime/primers/details?ensembl_version=81&primer_id=2013505) |
| TBP | TGCACAGGAGCCAAGAGTGAA | CACATCACAGCTCCCACCA | [2110939](http://bbcftools.epfl.ch/getprime/primers/details?ensembl_version=81&primer_id=2110939) |
| Gusb | GAAGTATCAGAAGCCCATTATTCAG | AACATCAGAGGTGGATCCTG | [1812772](http://bbcftools.epfl.ch/getprime/primers/details?ensembl_version=81&primer_id=1812772) |
| B2m | TGTACTACACTGAATTCACCC | CTTACATGTCTCGATCCCAC | [1932777](http://bbcftools.epfl.ch/getprime/primers/details?ensembl_version=81&primer_id=1932777) |
| A1AT | GACAGAAGGTCTGCCAGCTTA | TCAGCCCCATTGCTGAAGAC |  |
| ADH1B | CATGTGGCACAAGCGTCATC | CTTAAAGCCACCATAAACAGCC |  |
| CYP3A4 | GCCTGGTGCTCCTCTATCTA | GGCTGTTGACCATCATAAAAG |  |
| CYP3A7 | AAGTCTGGGGTATTTATGACT | CGCTGGTGAATGTTGGAGAC |  |
| CYP3A1 | TCCGGGACATCACAGACAGC | ACCCTGGGGTTCATCACCAA |  |
| CYPC219 | TGATCAAAATGGAGAAGGAAAAGC | TCTGTCCCAGCTCCAAGTAAG |  |
| CYP2C9 | TCCTATCATTGATTACTTCCCG | AACTGCAGTGTTTTCCAAGC |  |
| CAR | AAGTGCTTAGATGCTGGCATGAGG | CTCAGTTGCACAGGTGTTTGC |  |
| PXR | CCAGCCTGCTCATAGGTTCTT | AGCTACCTGTGATGCCGAAC |  |
| CEPBA | TATAGGCTGGGCTTCCCCTT | AGCTTTCTGGTGTGACTCGG |  |
| HLF | ATCAGACCAGGTCAGCTGTTG | GGCTCATAACCCACTGGGAC |  |
| CYP1A2 | CTTCGTAAACCAGTGGCAGG | AGGGCTTGTTAATGGCAGTG |  |
