## Supplementary material for "MIXL1 Activation in Endoderm Differentiation of Human Induced Pluripotent Stem Cells": ATACseq oligos

Supplementary Table 3: ATAC-seq oligo designs

| Ad1_noMix: | AATGATACGGCGACCACCGAGATCTACACTCGTCGGCAGCGTCAGATGTG |
| --- | --- |
| Ad2.1_TAAGGCGA | CAAGCAGAAGACGGCATACGAGATTCGCCTTAGTCTCGTGGGCTCGGAGATGT |
| Ad2.2_CGTACTAG | CAAGCAGAAGACGGCATACGAGATCTAGTACGGTCTCGTGGGCTCGGAGATGT |
| Ad2.3_AGGCAGAA | CAAGCAGAAGACGGCATACGAGATTTCTGCCTGTCTCGTGGGCTCGGAGATGT |
| Ad2.4_TCCTGAGC | CAAGCAGAAGACGGCATACGAGATGCTCAGGAGTCTCGTGGGCTCGGAGATGT |
| Ad2.5_GGACTCCT | CAAGCAGAAGACGGCATACGAGATAGGAGTCCGTCTCGTGGGCTCGGAGATGT |
| Ad2.6_TAGGCATG | CAAGCAGAAGACGGCATACGAGATCATGCCTAGTCTCGTGGGCTCGGAGATGT |
| Ad2.7_CTCTCTAC | CAAGCAGAAGACGGCATACGAGATGTAGAGAGGTCTCGTGGGCTCGGAGATGT |
| Ad2.8_CAGAGAGG | CAAGCAGAAGACGGCATACGAGATCCTCTCTGGTCTCGTGGGCTCGGAGATGT |
| Ad2.9_GCTACGCT | CAAGCAGAAGACGGCATACGAGATAGCGTAGCGTCTCGTGGGCTCGGAGATGT |

| **Tube #** | **Sample Name** | **Lane1** | **Lane 2** | **Barcoded Primer 2** |
| --- | --- | --- | --- | --- |
| 1 | FA2 A |  | x | Ad2.1 |
| 2 | FA2 B | x |  | Ad2.1 |
| 3 | FA2 C |  | x | Ad2.2 |
| 4 | FA3 A |  | x | Ad2.3 |
| 5 | FA3 B |  | x | Ad2.4 |
| 6 | FA3 C | x |  | Ad2.2 |
| 7 | MB2 A |  | x | Ad2.5 |
| 8 | MB2 B | x |  | Ad2.3 |
| 9 | MB2 C |  | x | Ad2.6 |
| 10 | FA1 A |  | x | Ad2.7 |
| 11 | FA1 B | x |  | Ad2.4 |
| 12 | FA1 C | x |  | Ad2.5 |
| 13 | MB1 A |  | x | Ad2.8 |
| 14 | MB1 B | x |  | Ad2.6 |
| 15 | MB1 C | x |  | Ad2.7 |
| 16 | MB3 A | x |  | Ad2.8 |
| 17 | MB3 B | x |  | Ad2.9 |
| 18 | MB3 C |  | x | Ad2.9 |
